# A whole-body cell type atlas mapped into an electron microscopy volume of an annelid worm

**DOI:** 10.1101/2025.10.30.685666

**Authors:** Oel Adam Phillip, Cros Cyril Christophe Daniel Robert, Pan Leslie, Buglakova Elena, Haury Parra Samuel, Santangeli Luca, Gerber Tobias, Papalopoulos Nikolaos, Vergara Hernando Martinez, Puga David, Disela Vanessa, Niederhaus Lara, Bertucci Paola Yanina, Witte Victoria, Bilgesu Kilic Asli, Quintana Urzainqui Idoia, Acevedo Julieta Maria, Musser Jacob, Kreshuk Anna, Arendt Detlev

## Abstract

Differential gene expression establishes the distinct physiology and morphology of cell types in an animal body. Single-cell sequencing and volume EM represent milestones toward the characterization of cell types, yet are difficult to combine for a comprehensive view on the cellular genotype-phenotype link. Here, we map a whole-body single-cell transcriptome into the *PlatyBrowser,* a multimodal cellular atlas for the marine annelid *Platynereis dumerilii*, and establish this combination uniquely for an entire animal. We learn that, in the 6-days-old worm, the majority of genes are tightly co-regulated to jointly implement one of eight major cellular morphotypes representing epidermis, gut, vasculature, myofibres, glia, motile cilia, glands, or neurons. Focusing on neurons, we uncover 14 families that by transcription factor identity, axonal projection, or sensory-secretory apparatus resemble conserved neuron types found in vertebrates, insects, or nematodes. We hypothesize that these existed in urbilaterian ancestors and represent the ancient core of nervous system centralization.

## Introduction

Building on the information stored in an animal’s genome, all cells that make an animal body express distinct subsets of available proteins to establish vastly different cellular morphologies and physiologies. Phenotypes are modular, with proteins and other constituents assembling into molecular machinery of different complexity. Relevant functions emerge at the level of single molecules such as enzymes; of molecular aggregates such as actin fibres, microtubuli, or vesicles; of complex composites such as pores, junctions, or synapses; and, finally, at the level of entire organelles such as the cilium.

Cell types represent a unique combination of molecular machinery, leading to distinct cellular phenotypes (Arendt, 2008). They are defined by their individual gene expression program that implements this machinery, under the control of a unique combination of transcription factors, referred to as terminal selectors^1–3^ that often assemble into Core Regulatory Complexes (CoRCs) (Arendt et al., 2015). Such individuated cell types represent evolutionary units that can be traced across species and across phyla ^4–7^. In recent years, rapid technical development has facilitated understanding cell type-specific differential gene expression and the resulting physiology and morphology.

First, the advent of single-cell transcriptomics and multi-omics has enabled the generation and comparison of cellular gene expression atlases, which reveal the transcriptomes of cell types^8–24^. To comprehensively capture and compare differential gene expression, however, such atlases need to comprise cells of the entire animal body, as available for human^25^, mouse^26^ or fly^17^. Full body atlases are more easily obtained for smaller animals such nematode, sponge ^27^, cnidarian^28,29^ ^30^, planarian^31,32^, or, more recently, the enigmatic *Xenoturbella*^33^. So far, however, these atlases are often incomplete in that tissues harbouring numerous cell types similar in morphology and/or physiology such as the nervous system are not fully resolved. Also, some atlases fail to distinguish developmental precursors from differentiating or fully differentiated cells. This blurs the recognition of cell types, because developmental precursors often have the mixed profiles of their later progeny. And since single-cell sequencing techniques deprive the cells of their morphological context, numerous spatial transcriptomics techniques have been developed based on imaging or sequencing^34^; yet, these efforts often cover subsets of cell types only and mostly lack cellular resolution.

Second, the past decade has seen immense progress in capturing cellular and sub-cellular morphologies via the rapid advancement of volume EM techniques, such as serial block-face^35^ or focused ion-beam-milling^36^ scanning electron microscopy (FIB-SEM and SBEM). This goes in concert with advanced tracing techniques and with deep-learning based automated segmentation pipelines for cellular shapes, organelles, or nuclei^37–39^. Again, however, to comprehensively capture and relate all the distinct cellular morphologies that belong to an animal, these volumes need to span the entire body, which can only be achieved for small animals ^27–33^. In addition, fixation and processing for EM is yet incompatible with sequencing efforts, obscuring the link between the imaged tissue and cells of interest and the underlying differential gene expression.

In a first effort to establish a link between gene expression and volume EM, we have recently created the *PlatyBrowser*, an interactive tool to explore multimodal datasets mapped on a high-resolution serial block-face EM volume of a 6-days-post-fertilization (6dpf) young worm of the marine annelid *Platynereis dumerilii* ^37^. Up to this stage, *Platynereis* development is highly stereotypic, which allows the registration of different volumes of diverse modalities.

Leveraging this, we have mapped the whole-body expression patterns for more than 250 differentially expressed genes, obtained through “Profiling via Signal Probability Mapping” (ProSPr)^40^, to the EM volume. Furthermore, we have traced the axons for 500 brain neurons and used automated segmentation to capture cellular shapes, nuclei, and chromatin, which we use to define so-called “morphofeatures” via deep learning for each cell^41^.

Leveraging and expanding on this resource, we now add a full whole-body single-cell transcriptome to the *PlatyBrowser -* creating a link between cellular transcriptomes and (sub-) cellular morphology for an entire body. Sequencing 100,000 captured nuclei for the entire 6dpf *Platynereis* young worm, we use repeated sub-clustering and merging of cellular clusters, combined with manual curation, to identify around 170 genetically individuated cell types that make up the worm body. We then use the differential expression of ProSPr-genes to map these cell types to the *PlatyBrowser,* complemented by a newly developed pipeline for the mapping of additional cell type-specific marker genes that we use to refine and validate the localisation of cell types into the new *PlatyBrowser*, which is now publicly available under <u>URL</u>.

Leveraging this new resource, we reveal the existence of huge sets of genes that are tightly co-regulated across defined subsets of cell types, which we refer to as “differentiation coregulons”. We find that genes that belong to the same differentiation coregulon jointly encode complex cellular features that are characteristic for epidermis, muscle cells, gut cells, motile ciliated cells, glia cells, macrophages, gland cells, and neurons; and they show shared transcription factor motifs on accessible DNA as revealed by bulk ATAC-sequencing. Focusing on neurons, we show that these belong to 14 neuronal families that share specific combinations of transcription factors. With our new resource we co-characterize these families molecularly and morphologically, regarding axonal projection, transmitter usage, secretion, sensory processes with or without cilia, and receptor usage. Strikingly, most of these match vertebrate neuronal families by both terminal selector code and specific molecular and morphological features - and thus likely existed in urbilaterian ancestors. The *Mnx+* cholinergic motor neurons and the *Evx+* and the GABAergic *Dbx, Ptf1a+* commissural interneurons emerge as candidates for ancient central pattern generators in the trunk. In the brain, *Dbx+, Ptf1a+* interneurons and the closely related *Ptf1a, Arx+* mushroom body neurons match telencephalic neuronal families, which thus qualify as possible evolutionary precursors for sensory associative centres. We identify several families of mechanosensory neurons, among which the *Emx+* neurons populate the palpae, which are appendages for food testing; and the dualism of somatic *POUIV+* mechanosensory and autonomic *Phox2+* proprioceptive neurons likewise represents ancient heritage. Beyond that, *Rx+*, *Bsx+* and other sensory-neurosecretory, hypothalamus- and placode-like cells combine ambient light detection via OPN3, OPN5, RRH, and Go-opsin and chemical sensing via vertebrate-type olfactory receptors, representing the main sensory input into the brain. Building on this, we present a first reconstruction of the urbilaterian nervous system as a working hypothesis and first step towards understanding nervous system centralisation.

## Results

### Single-cell sequencing identifies individuated cell types

We performed single-nuclei RNA-seq across 6 different larval batches (14 10X Genomics libraries) and obtained 92136 nuclei after quality control and doublet removal (Figure 1A). This number is 10 times more than the average number of cells of a Platynereis larva at 6 dpf providing enough coverage that we likely sampled each cellular pair (bilateral homologs) in the body. Using Seurat’s default parameters (resolution 0.8), we obtained 65 clusters (Figure 1B). This did not fully resolve known diversity as distinct cell classes (e.g., muscle and neurons) were clustered together masking internal heterogeneity. To address this, we generated a neighbor-joining tree and performed a bootstrap analysis on the tree which allowed us to assign probability scores to each of the nodes (Figure 1C). This resulted in 13 clades for further analysis of which clade 3 was split into two well-supported subgroups. 11 standalone clusters were assigned as “no clade”. Marker gene searches revealed canonical markers for each of the clades allowing us to assign broad labels to the clades (Supplementary Figure 1B).

**Figure 1.**
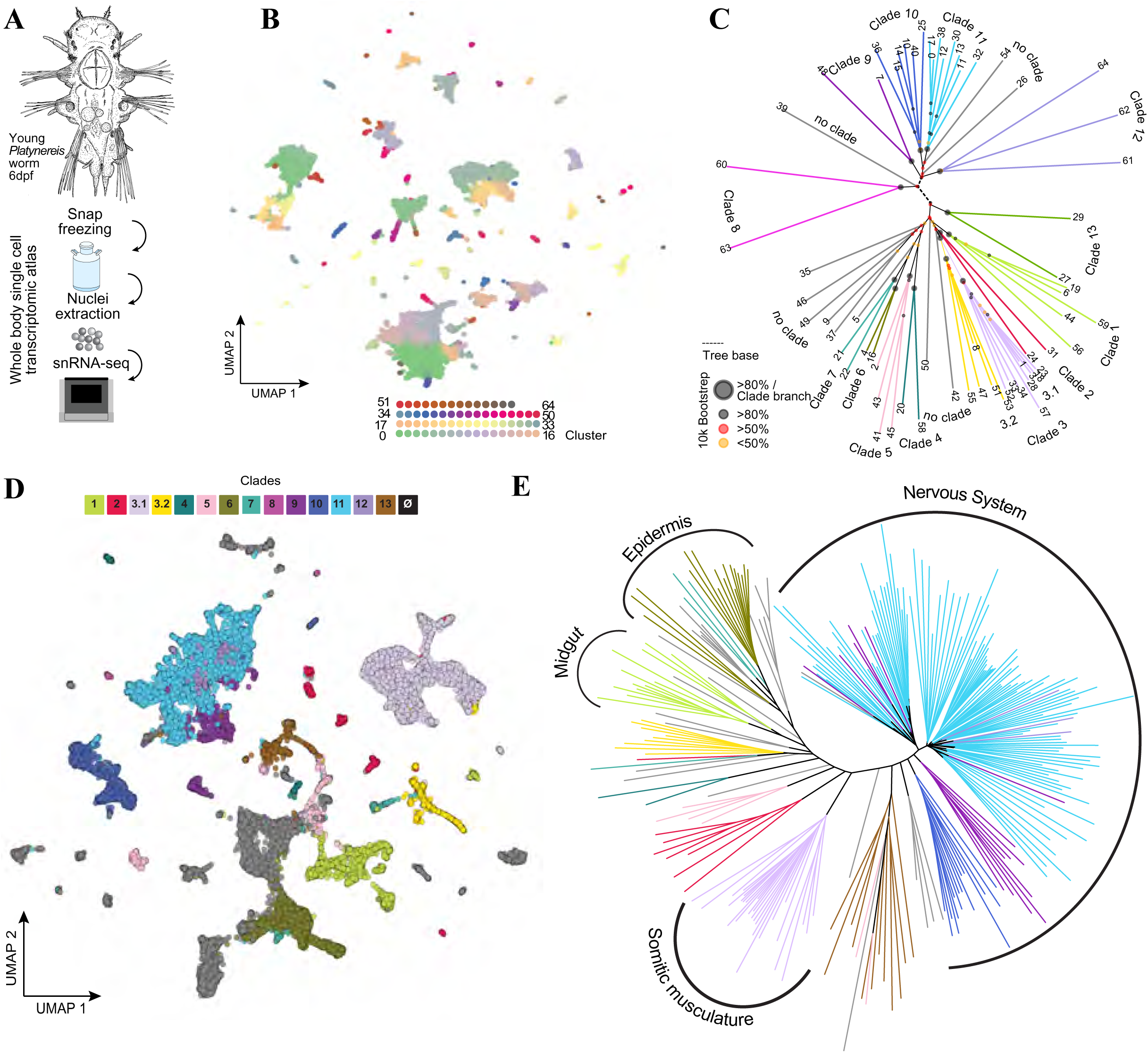
Analysis of the 6dpf single nucleus object. **A)** Schematic representing the experimental workflow for the snRNA-seq atlas generation. **B)** UMAP embedding contains all nuclei of the atlas grouped across 65 color-coded Louvain clusters. **C)** Neighbor joining tree with bootstrap values for each node visualizes the grouping of clusters into clades. Bootstrap values of more than 80% were considered for cluster groupings. **D)** Only differentiated cells based on the clade subclustering were embedded in a UMAP. Original clade identities are shown on the embedding. **E)** Neighbor joining tree of terminally differentiated cell types is shown. Clade assignment reveals initially wrongly assigned cell types. Cell type labels based on canonical markers are indicated.

The Platynereis 6dpf young worm already comprises many differentiated cells but also developmental precursors that often show generic expression profiles and mask the more specific transcriptomic profiles of the differentiated cell types. Since our main objective was to obtain a close-to-complete set of differentiated cell types for the 6dpf young worm, we needed to distinguish developmental precursors from differentiated cells. To this end,we subclustered each of the clades and screened for subclusters exhibiting a rich catalog of specific gene expression, indicative of “individuated” differentiated cell types. Developmental precursors with very few or no specific markers were no longer considered. To improve resolution, several of the obtained subclusters had to be subclustered again. This manual inspection allowed us to identify maximally homogenous units with unique gene expression profiles, representing individuated cell types. Individuation was apparent at the level of terminal selector transcription factors and effector genes, indicative of cell type-specifc expression programmes. After cleaning the data from non-individuated subclusters and subsubclusters, we obtained an atlas of 31,121 fully differentiated cells across 173 cell types. These curated individuated cell types were tagged with a unique identifier comprising their initial clade and subcluster of origin (cladeXsubY); and subsubcluster if a second subclustering had been performed (CladeXsubYsubsubZ). We next generated from this new data set a Neighbour-joining tree which now represents a cell type tree (Figure 1D+E) in contrast to the developmental tree in Figure 1B. Remarkably, this second tree shows new clades that roughly correspond to the old clades; yet we observe many cell types that change clades. This illustrates the difference between a developmental and an evolutionary cell type tree; in essence, cell types with dissimilar differentiation profiles may develop from the same developmental precursor (and developmental trajectory), and cell types with similar (evolutionarily related) differentiation profiles may develop from distinct precursors.

### Mapping cell types to the EM volume in *PlatyBrowser*

For the previous version of the *PlatyBrowser*, the expression of around 250 genes had been registered to the EM volume via Profiling by Signal Probability mapping (ProSPr)^37^, which conveys each segmented cell in the EM volume with a characteristic molecular fingerprint that can be used for the mapping of cell types that express similar combinations of genes. Leveraging the differential co-expression of ProSPr genes, we set out to map the curated individuated cell types from the 6dpf scRNAseq object. To streamline this process, we established a pipeline for the registration of 3D volumes for the expression of missing marker genes to *PlatyBrowser*, using HCR-based FISH and two different imaging modalities (Fig. 2A). The registered expression is visualized in the new edition of the *Platy*Browser under <u>URL</u> and the genes listed in Suppl. Fig. 2. We tested the accuracy of image registration for genes expressed in morphologically distinct structures, as exemplified for *r-opsin* expression in the adult eyes (Fig. 2B). This is a versatile pipeline for the mapping of any gene of interest in the future.

**Figure 2.**
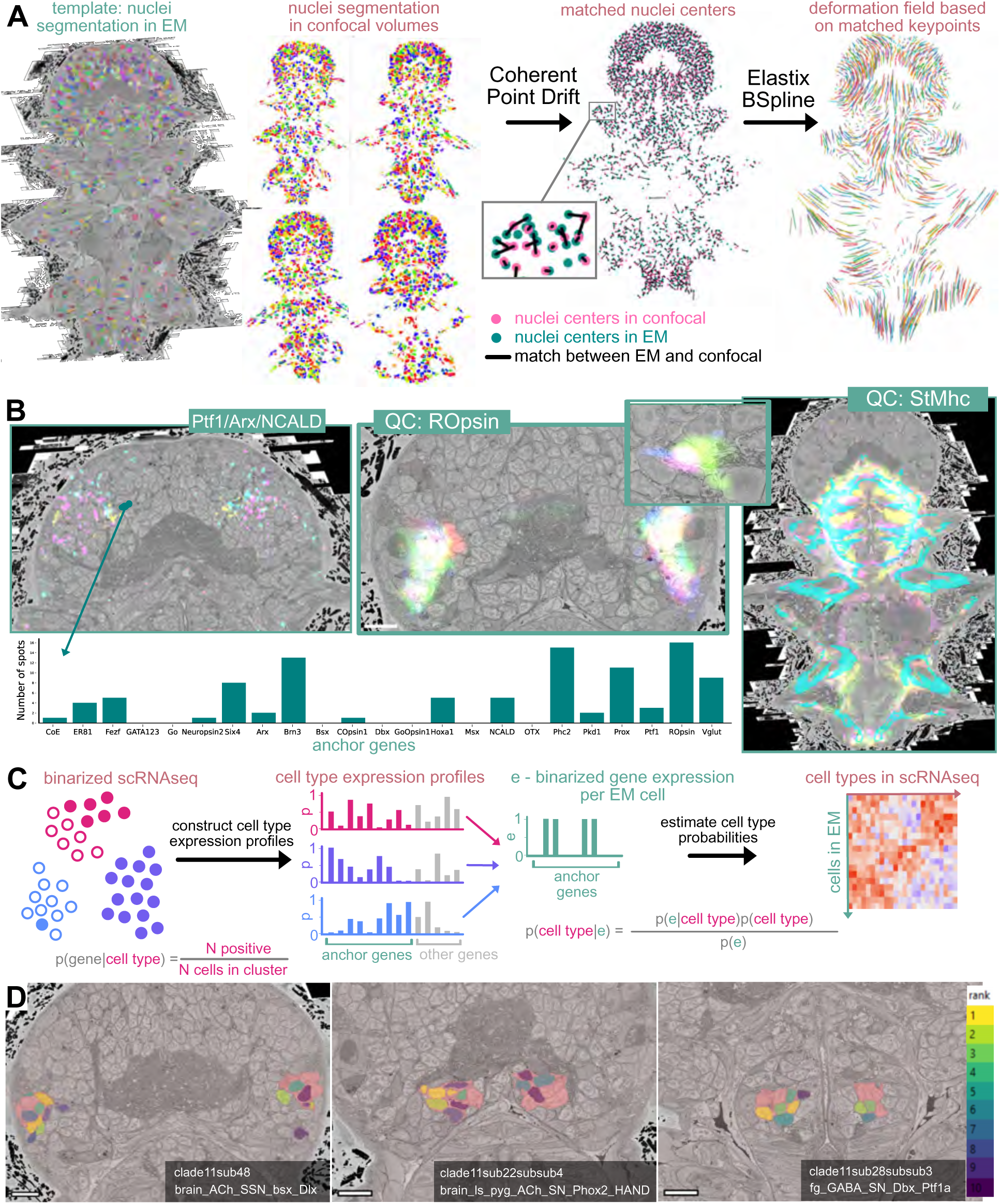
Analysis of the 6dpf single nucleus object. **A)** General workflow of registration pipeline. From left to right: Reference EM volume nuclei segmentation, examples of nuclei segmentation of confocal imaged DAPI channel showing the difference in nuclei location between specimens, example coherent point drift matching of EM and confocal nuclei, example Elastix BSpline confocal volume deformable alignment based on matched EM nuclei. **B)** Quality control visualizations of registration pipeline. From left to right: HCR spot registration of known MB markers (Ptf1 - blue, Arx - magenta, NCALD - yellow), validation of pipeline accuracy through known eye-cell marker ROpsin with different specimen HCR stainings in different colors, validation of pipeline accuracy through comparison of known muscle marker StMhc of different specimen HCR stainings in different colors and previous ProSPr registration in cyan. **C)** Schematic representation of the automated cell type annotation pipeline. **D)** Visualization of automatic annotation results of selected cell types as well as independent manual annotation for reference. Automatically annotated probability colored by ranking of cell type in the overall probability assignment (yellow-to-blue). Rank-1 indicates the cell type being most probable, rank-10 the 10th most probable. Independent manual annotation, taking into account bilateral symmetry and additional specific markers, colored in red. 5 micrometer scale bar for reference.

We then created an automated pipeline that maps the curated cell types to the segmented EM nuclei in *PlatyBrowser.* This was achieved by inferring the ProSPr and image registration expression profiles for each nucleus and systematically comparing these to the snRNA-seq cell type expression profiles. The comparison was performed by calculating the likelihood of a nucleus belonging to a given cell type based on a specificity-based weighted matching of snRNA-seq and ProSPr/imaging gene expression (Fig. 2C). Having calculated this probability for all cell types, we displayed the nuclei that belonged to a cell type of interest with highest probability on *PlatyBrowser*. This was followed by manual curation for each cell type, taking into account bilateral symmetry and known gaps for ProSPr patterns (Fig. 2D). Automatically located and curated cell types are visualized in the new version of the *Platy*Browser (<u>URL</u>), and as exemplified in Figure 2E-H. The combined morphological and molecular information achieved through the mapping allowed us to assign meaningful names to cell types in the format “Bodypart_transmitter_modality_CoRC” (Fig 2D). We validated the mapping of selected cell types through the registration of expression volumes for highly cell type-specific genes available in *PlatyBrowser*. For example, the Trp channel Pkd1.1 is only expressed in clade10sub3subsub5-9 (brain_or_ls_pyg_Glu_mechSN_POUVI_Lhx3). For localisation validation we also took advantage of cell type-specific features that we could visualise in both light and electron microscopy, such as traced commissures (see below).

### Large sets of genes coregulated across cell types

Since the neighbour-joining tree of curated differentiated cell types based on entire cellular transcriptomes (Fig. 1D) revealed strongly supported clades of cell types, we investigated the differential expression of genes supporting these clades across the individuated cell types. We followed two strategies to identify such gene modules. First we swapped our raw count matrix of the data set containing only differentiated cells in order to cluster genes and not cells (Supp Figure 3) revealing 42 gene clusters. When we scored our atlas for these gene modules an association of clusters to individual but also across clades was found (Supp Figure 3C-D). Corroborating this, a weighted gene co-expression network analysis (WGCNA) revealed large sets of genes uniquely and consistently active across clades that were broadly overlapping with the gene clusters above (Fig. 3A+B; Supplementary Figure 3F+G; Supplementary Figure 4A+B). These findings indicate the existence of large sets of coregulated genes in the *Platynereis* genome, which we refer to as ‘differentiation coregulons’.

**Figure 3.**
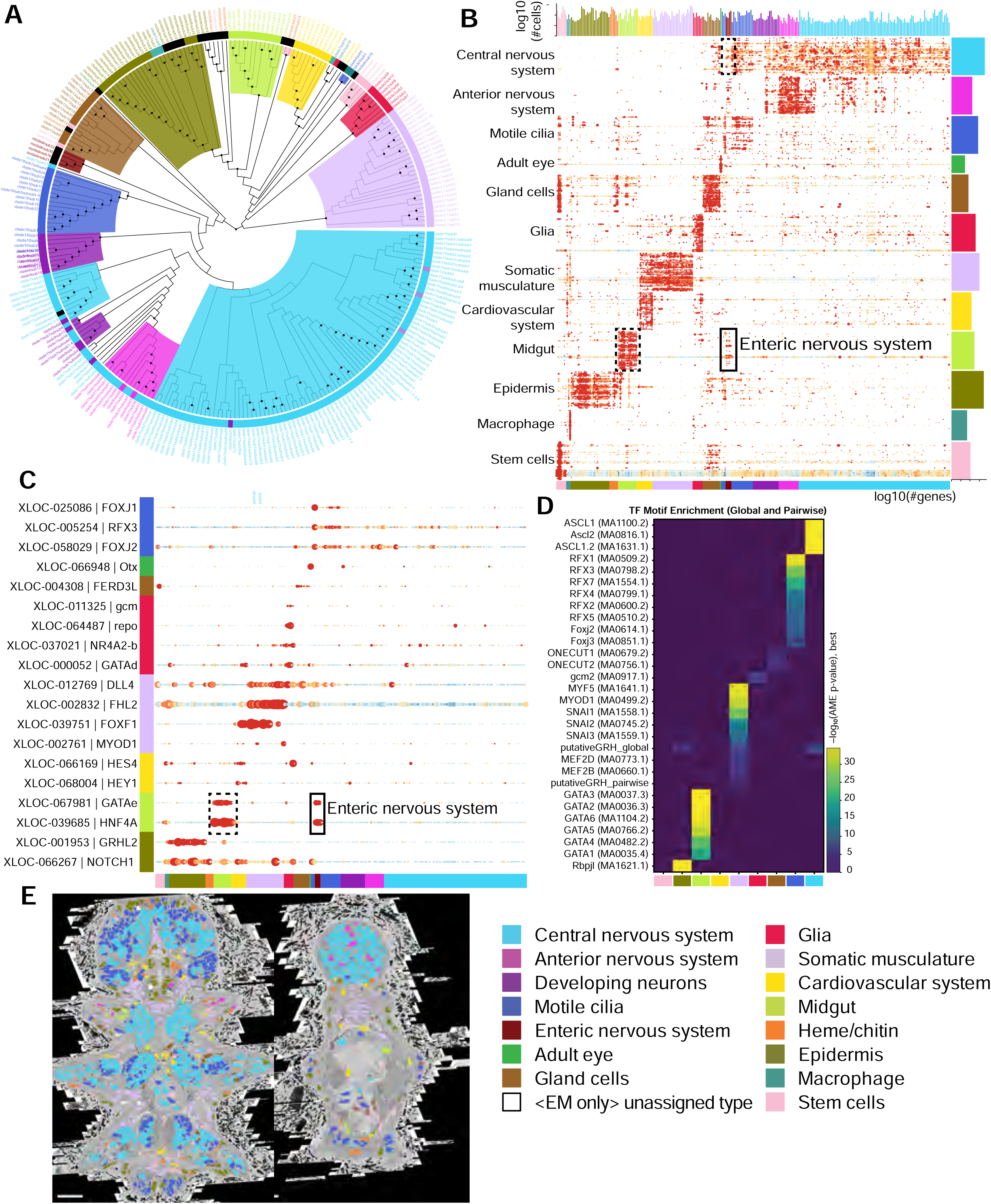
Differentiation coregulons. **A)** WGCNA coregulons mapped onto the cell type tree introduced in Figure 1E. Color code as in B). The outer periphery of the tree represents the initial clade origin for individual cell types as in Figure 1B+C. **B)** Expression for the 60 most representative genes for each coregulon plotted against all curated differentiated cell types. Note that coregulon expressions are partially overlapping, in particular with the expression of the neuronal coregulon. **C)** Transcription factors specific to individual coregulons. **D)** Heatmap showing the enrichment of transcription factor binding motifs across major coregulon modules in ATAC-seq peaks. Motif enrichment significance (−log_10_ p) was estimated using AME, and the putative *de novo* Grainyhead (GRH) motif identified by TOMTOM comparison was appended for reference. The observed motif enrichment patterns together with the expression profiles of the corresponding TF support the assignment of cell types to coregulons. **E)** Mapping of the coregulons into the *PlatyBrowser*.

Most of these genes were effector genes driving clade-specific differentiation. Gene ontology (GO) term analysis revealed candidate functions for these genes, suggesting that they were jointly encoding clade-specific cellular morphology and architecture. To validate this, we inspected the tissue contribution, morphology, and architecture of the cell types in which a given coregulon was expressed, taking advantage of the *PlatyBrowser* mapping of partaking cells (Fig. 3D). Taking into account genes and morphology, our inspection revealed coregulons demarcating epidermis, macrophage-like cells, midgut, visceral musculature, somatic myofibres, motile ciliated cells, glia cells, gland cells, and neurons.

Importantly, we also identify coregulon-specific transcription factors, such as Grainyhead for epidermis, GATAe and HNF4 for midgut, Hes4 and HEY for visceral musculature, MyoD and FoxF1 for somatic musculature, FoxJ and Rfx1/2/3 for motile ciliated cells, Gcm and Repo for glia cells, Ferd3l for gland cells, and Myt1l for neurons. These factors are exclusively and consistently expressed in their coregulons and thus identify as prime candidates for coregulon-specific gene regulation. Notably, many of these transcription factors are already known to specify similar tissues and cell types in other bilaterian groups, suggesting that these tissues, together with their specifying transcription factors, are evolutionarily ancient (see discussion).

To investigate whether these transcription factors indeed control coregulon-specific gene expression, we performed ATACseq to probe motif grammar of chromatin accessible regions. Motif enrichment analysis of coregulon associated peaks reveal that each coregulon displays a characteristic but partially overlapping motif signature (Fig. 3D). Moreover, we were able to identify de novo motifs for genes of the epidermal coregulon that may bind Grainyhead, due to their palindromic nature and similarity to known Grainyhead motifs. While some motifs are only enriched only in one coregulon such as FoxJ and Rfx1/2/3, others are found across different coregulons. However, the restrictive expression of the corresponding transcription factors suggests that only a subset of the motifs are functionally engaged in any given tissue. The modular combination of shared cis-regulatory grammar and tissue-restricted TF activity likely underpins cell type specification and evolution.

Searching for transcription factors specific for neuron types, we find the *Myt1l* gene expressed in all neurons and gland cells, which indicates that neuronal identity might depend on the repression of non-neural differentiation programmes^42^. All neuronal types were strongly enriched for homeodomain binding sites, in line with the widespread expression of homeodomain transcription factors (Fig. 3x). In particular, all neuronal cell types show combinatorial expression of at least one member of the LIM, POU, and Cux homeodomain families (Suppl. Fig. 4C) Interestingly, we find a small group of enteric neurons that represent an overlap of both, midgut and neuronal coregulons. These neuronal cell types show the signatures of endocrine pancreas (mnx, lmx1, nkx2.2 for nocladesub16 and nocladesub21) versus exocrine pancreas (Ptf1a for nocladesub32). We also observe neuron types sharing the neuronal and the motile cilia coregulon (Fig. 3B).

### The combinatorial expression of transcription factors defines neuronal families

To explore and compare the distinct identities of neuronal cell types, we focused on highly differentially transcription factors specific for subsets of the neuronal cell types, many of which represented so-called terminal selector genes^43^. We identified around 80 transcription factors with partially overlapping expressions in a total of around 110 neuron types (Figure 4A; Supplementary Figure 5). About half of these factors are homeodomain factors mostly of the NK and Paired families, followed by basic Helix-loop-helix factors and nuclear receptors. Given the importance of many of these factors in specifying neuronal fates, we propose that the groups of cells showing similar combinations of transcription factor expression represent families of neuron types that are evolutionarily related (neuronal families, see discussion).

**Figure 4.**
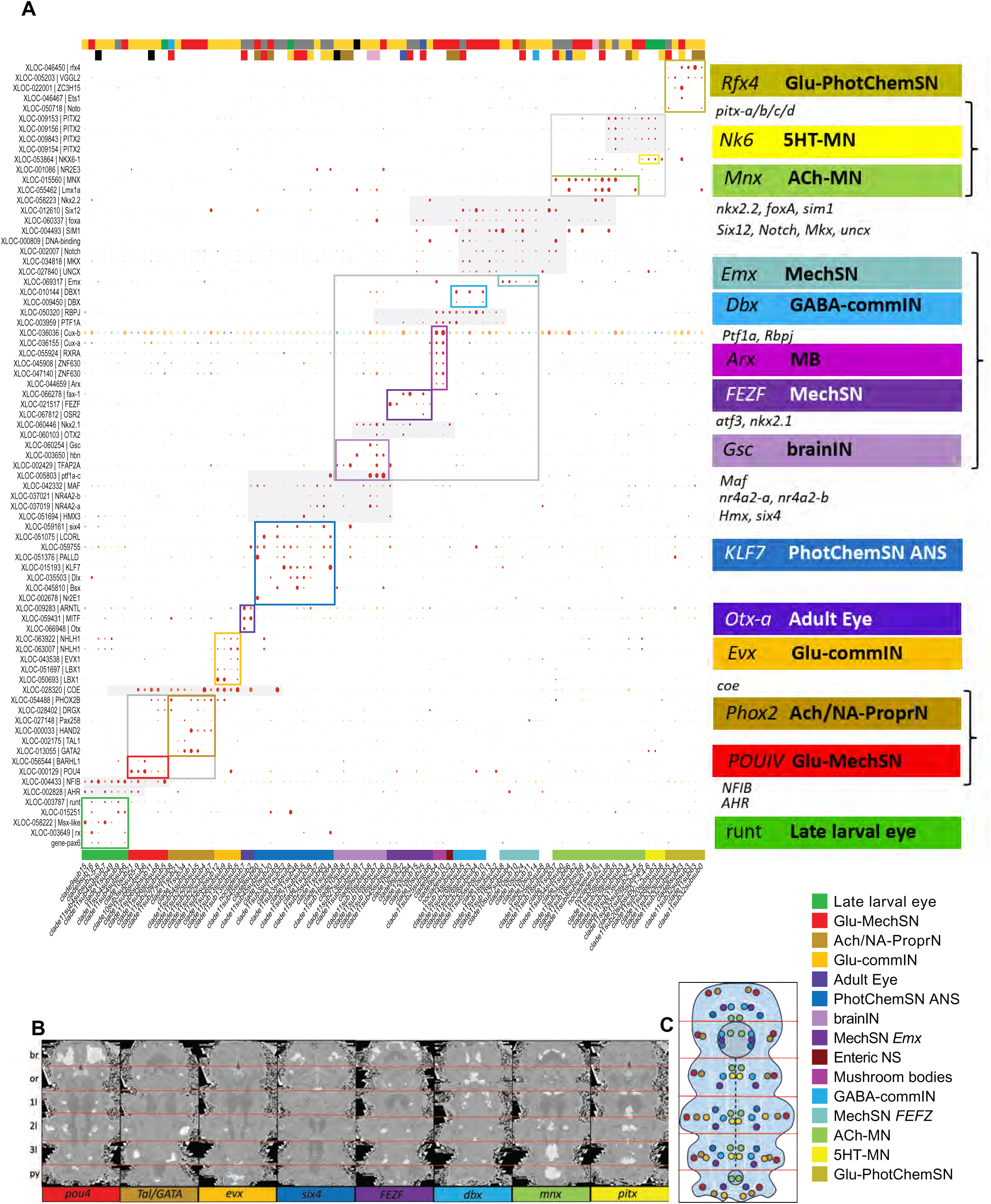
Transcription factor-defined neuronal families. **A)** Differentially expressed transcription factors plotted against curated neuronal types. Colored boxes indicate neuronal families. Grey boxes represent superfamilies. Transmitter types and the presence of mechano-, warmth, light and chemoreceptors indicated below the neuronal types. Family-specific transcription factors and characteristics as determined by combined molecular and morphological analysis indicated on the right. See Supplementary Figure xx for a larger version of this panel with gene and cell type names. **B)** Mapping of neuronal families into the *PlatyBrowser* via the expression of family-specific transcription factors **C)** Summary scheme for the presence/absence of neuronal families in the six body segments of the *Platynereis* young worm. Colour code as in A. Note that while most neuronal families are represented in all segments, some inter- and all motoneuron types are missing from the nervous system in the oral and pygidial segment and instead move into the fore- and hindgut anlage. Segments: br, brain, or, oral; 1l, 2l, 3l, 1st, 2nd, 3rd larval; py, pygidial.

Notably, POU and LIM homeodomain family members are expressed more broadly across neuronal types (Supplementary Figure 4C) and thus mostly not included in Figure 4A. Plotting the differentially expressed transcription factors subdivides the full complement of curated neuron types into 14 groups, each of which specifically co-expresses several transcription factors (Figure 4A). Given the cross-family expression of some factors, these families can be grouped into three overarching groups (grey boxes in Figure 4A). The first (medial) group contains motor neuron families co-expressing *Nkx6* and found in the medial parts of the CNS; the second (intermediate) group has mixed interneuron and sensory types as well as families restricted to the brain co-expressing co-expressing Ptf1a and Rbpj (see below); the third (lateral) group contains lateral and peripheral sensory neuron types co-expressing *COE* and *Phox2B*.

To molecularly characterise the neuronal families, we plotted the expression of genes for transmitter synthesis and release (Supplementary Figure 6+7), genes encoding mechanosensory and warmth-sensitive Trp channels (Supplementary Figure 8), various opsins (Supplementary Figure 9), genes encoding chemosensory and olfactory G-protein coupled receptors (Supplementary Figure 10), and genes encoding neuropeptides (Supplementary Figure 11) against cell types and found various family-specific patterns, corroborating our family subdivision. This allowed us to name the families according to transmitter usage and sensory modality as listed in Figure 4A.

Mapping these families to the *PlatyBrowser* via the expression of family-specific transcription factors (Figure 4B), we find that most neuronal families are found across all six body segments of the young worm, i.e. brain, oral, 1st, 2nd and 3rd parapodial, and pygidial as serial homologs (Figure 4C). Within these segments, they locate to different mediolateral positions. In line with this, the family-specific transcription factors include conserved mediolateral patterning genes of the NK and PAX homeodomain transcription factor families (e.g., nkx2.2, nkx6, pax6, pax258). Notably, only neuronal families that belong to the lateral group have neuron types present in the CNS and PNS of all six body segments. Neuronal families that belong to the medial and intermediate family are lacking CNS representatives in the oral and in the pygidial segments. Instead, the serial homologs of the oral and pygidial segments are internalized and form part of the foregut and hind-midgut, respectively (Figure 4C).

### Identification of motor neurons and of GABAergic commissural interneurons

Our new resource uniquely enables the in-depth characterisation of neuron types representative of the neuronal families via the correlation of expression profiles with available cell shapes, cellular ultrastructure, and axonal tracings.

The homeodomain factor Mnx is a conserved motor neuron marker across phyla^44^. We found *Mnx+* neurons in the brain, the foregut, and the three larval segments. Previous studies had revealed a prominent pair of brain motoneurons in the *Patynereis* young worm with contralateral descending axons directly innervating the longitudinal muscles, which play a role in bending the body during phototaxis. We validated the identity and projection of the *Mnx+* brain neurons via F0 transgenesis in 6dpf worms injected with an Mnx promoter construct. The tested construct includes 4174 bp of the Mnx genomic regulatory region immediately upstream of the ATG, driving the expression of a fusion protein consisting of tdTomato with three hemagglutinin (HA) tags and an artificial palmitoylation site (Palm-3xHA-tdTomato). The construct also expresses H2B-EGFP under the control of the ubiquitous promoter of the P2 RNA-binding protein^45^ (Fig. 5A) and found it to be expressed in the brain motor neurons, with contralateral innervation of the longitudinal muscles (Fig. 5A’). Furthermore, we found that the cholinergic and glutamatergic *Mnx+* and *Pitx+* motor neurons of the trunk share expression of the protostome-specific neuropeptide myomodulin^46^, known to modulate ion channels in the innervated musculature^47^. This profile is shared by the two enteric neuron types (nocladesub16, nocladesub21), which besides *Nkx2.2* express the parahox transcription factors *Pdx* and *Cdx* and thus exhibit the conserved molecular signature of endocrine and exocrine pancreatic cells and of gut cells and neurons in vertebrates and in the sea urchin^22^. Besides myomodulin-producing cells, the *Pitx+* family contains neuron types with the full complement of enzymes, transporters, and channels for 5HT release and re-uptake, indicative of serotoninergic neurons.

**Figure 5.**
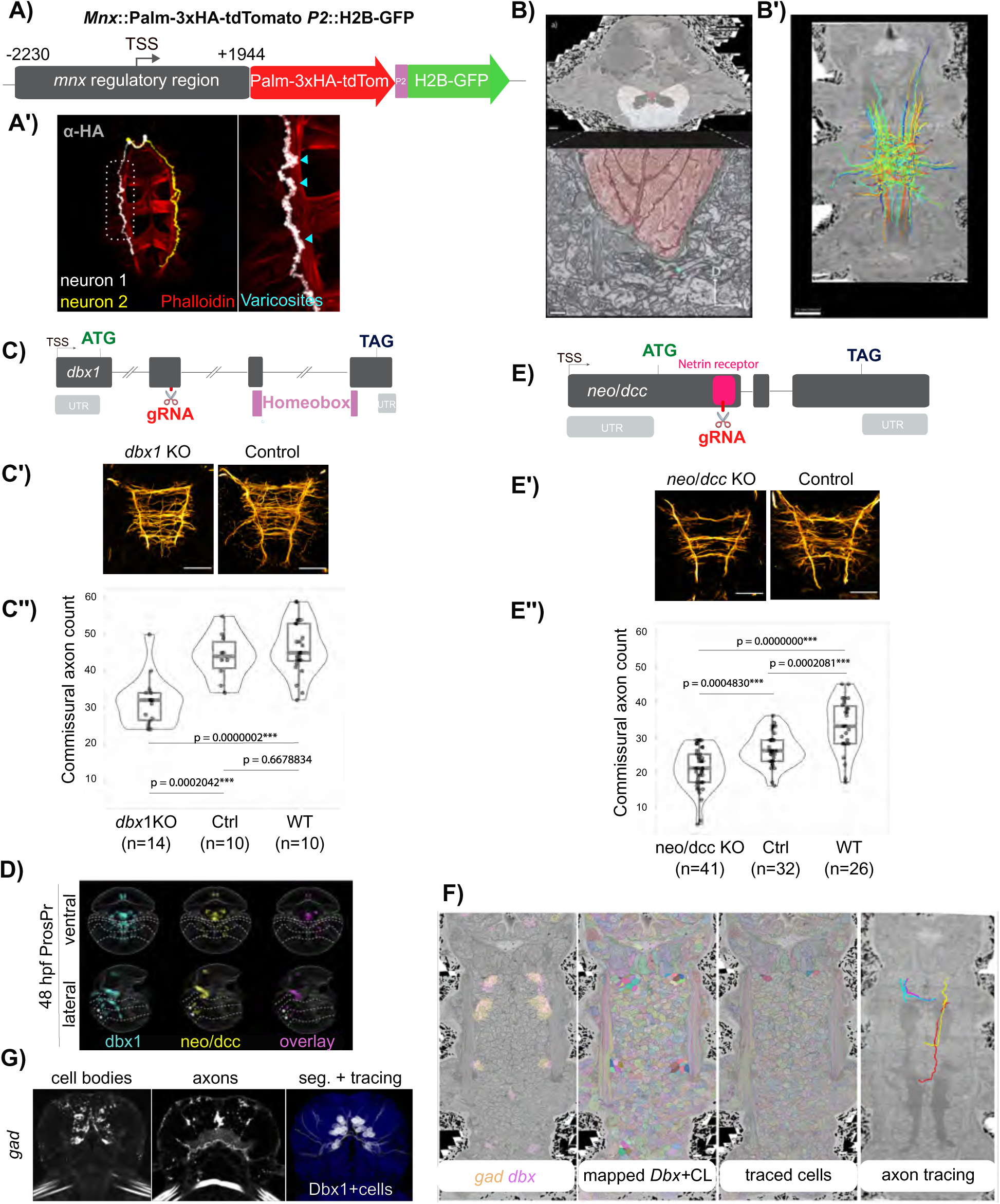
Commissural neurons in the 6dpf trunk. **A)** Schematic representation of the *mnx* reporter construct **A’)** Left panel: 6dpf animal injected with the *mnx* reporter construct and stained with anti-HA antibody and phalloidin (red). The HA positive signal of two neurons (white and yellow) is shown. Right panel: magnified area marked demarcated in the left panel showing varicosities (arrowheads). **B)** Commissural neuron tracing on the 6 dpf SBF-SEM volume. Upper panel: Cross section at the level of the second segment. VNC soma (white). Axochord in red. VNC neuropil (electron dense dark grey area, not coloured). Lower panel: Neuropil and axochord at higher magnification. Individual axons can be distinguished. A seeding point (cyan asterisk) was set in every axon of the second segment that we could see crossing from one side of the axochord to the other. (Scale bar in upper panel: 10 um and lower panel: 0.5 um). **B’)** Ventral view of the traces of the commissural neurons, each in a different colour. A single plane of the SBF-SEM dataset is visible, but belongs to a slightly more dorsal position so as not to interfere with the commissural traces. The segmental boundaries are indicated with dashed lines. **C-E)** Schematic representation of the **(C)** *dbx1* and **(E)** *neo/dcc* genomic locus indicating the target site of the gRNA utilised to generate the CRISPR KOs. **C’-E’)** Maximum projections of confocal stacks of **(C’)** *dbx1* KO and injected control (D’) *neo/dcc* KO and injected control. Representative images are shown. **C’’-E’’)** Commissural axon count for the different genotype larvae stained with an antibody against acetylated tubulin. Each point represents the number of axons counted in one larva. (n) number of larvae quantified for each genotype. The p values are from Tukey’s Test after ANOVA. Sig. * < 0.5, ** < 0.1, *** < 0.01. **D)** Expression patterns of *dbx1* and *neo/dcc* at 48 hpf are shown in the left and middle panels. The right panel shows the overlay expression of the two genes. The dashed delineate the anterior boundaries of segments 1, 2, 3, and 4. The asterisks show expression domains where the expression patterns of *dbx1* and *neo/dcc* overlap. **F)** 1st panel: PlatyBrowser view showing co-expression of *gad* and *dbx1+* in the trunk of a 6dpf worm. 2nd panel: Localisation of Clade11sub5subsub10 (ls_GABA_cSN_Dbx_Ptf1a) to the ventral nerve cord. 3rd: Traced cells of Clade11sub5subsub10. 4th: Traced axons for these cells in the 1st larval segment. **G)** Expression of *gad mRNA* in the 6dpf brain detected by HCR staining. Left panel: cell body expression, middle panel: axon expression. right panel: manual single-slice segmentation and axonal tracing of Dbx1+ commissural brain cells.

To identify commissural neurons in the ventral nerve cord, we traced all axons passing contralaterally below the axochord^48^ in the second larval segment (Figure 5B upper panel) and identified the corresponding cell bodies. We distinguished ascending and descending axons (Fig. 5B lower panel). The full complement of traced axons is shown in Fig. 5B’. We assigned expression profiles to the identified commissural cells and identified 8 commissural neuron types including *Phox2+*, *Dbx+, Evx+*, and *Pitx+* neurons. To validate the role of the Dbx transcription factor in the specification of commissural neurons, we injected guide RNAs targeting an off target free region in the second exon of Dbx (Fig. 5C) together with CRISPR-Cas9 mRNA into one-cell-stage embryos and thus produced F0 animals with partial or full knockout of the Dbx gene. In the F0 Dbx knockout animals, we found the number of commissures significantly reduced (Figure 5 C’ and C’’). We also observed that neogenin, a DCC receptor for netrin, is specifically expressed in *Dbx+* cells from 48 hpf onwards (Fig. 5D). We accordingly produced neogening F0 KO animals using a gRNA targeting the netrin receptor region (Figure 5E) which causes a disruptive deletion and found the number of commissures significantly reduced in the Neogenin F0 CRISPants compared to controls (Fig. 5E’ and E’’). These findings strengthen the notion that Dbx is required for the specification of commissural neuron types that deploy the Netrin/neogenin ligand receptor system for commissural outgrowth. Corroborating this, we also traced the axons of the *Dbx+* neurons in the 1st larval segment and confirmed their commissural nature (Fig. 5F).

The *Dbx+* neuron types express enzymes and transporters for GABAergic transmission, as validated *in situ* by the specific co-expression of *Dbx* and *Glutamic Acid Decarboxylase (GAD)* (Fig. 5F). They belong to a larger neuronal family that specifically co-express the Ptf1a/RBPJ transcription factor complex, which is known for its widespread role in the specification of inhibitory GABAergic interneuron types in the vertebrate neural tube^49^. *Dbx+, Ptf1a+, RBPJ+* GABAergic neurons were also found in the brain, and their commissural nature validated via HCR staining of *GAD* transcripts, which were detected in cell bodies and outgrowing axons including brain commissures (Fig. 5G). In *Platynereis*, the larger Dbx, Ptf1a neuronal family also contains the two neuron types representing the mushroom bodies (Fig. 4A). Finally, we also validated the commissural nature of *Evx+* neuron types in the 3rd larval segment via the targeted tracing of neurons in the EM volume to which these cell types were mapped (data not shown).

### Molecular-morphological characterisation of sensory neurons

Mechano-, photo-, and -chemosensory neurons were identified through expression of Trp (Transient receptor potential) or Piezo channels, opsins, or various conserved olfactory receptors (see above). We found detection of light and of chemicals often occurring in the same neurons, so that we broadly distinguish mechano- versus photo-/chemosensory neurons (Supplementary. Figure. 8).

Mechanosensory neurons express the motile cilia coregulon that includes Trp channels and other proteins implicated in the formation and function of hair cells in the vertebrates, such as NOMPC^50^, the stereocilia linker BAIAP2^51^, or the stereocilia coat protein Pkhd1l1^52^. Exemplifying this, the *Emx+* family of mechanosensory neurons specifically expresses the motile cilia coregulon with NOMPC, known to convey a sense of gentle touch (Yan et al 2013). We located these *Emx+* neurons (e.g., clade10sub11; Fig. 6A) to the palpae, a pair of head appendages involved in food sensing via sensory cilia, which thus identify as mechanosensory. In line with this, NOMPC channels are involved in tactile food sensing in *Drosophila* ^53^and *Caenorhabditis elegans*^54^. The *Emx+* family also contains two dopaminergic (DA) mechanosensory neuron types co-expressing tyrosine hydroxylase (TH), dopamine transporter (DAT) and the vesicular monoamine transporter (VMAT), as well as the the aryl-hydrocarbon receptor (AHR), which binds dopamine (ref). One of the DA neuron types (clade10sub7subsub4) is found at the base of the palpae, close to the palpal nerve, and another next to the stalk of the mushroom bodies (Fig. 6B). The *Emx+* DA neurons send out highly bifurcating short axonal projections and long branching sensory processes. They also express the receptor for non-noxious warmth, TRPM2^55^, and the thermosensitive TrpV3 (Supplementary. Figure. 8).

**Figure 6.**
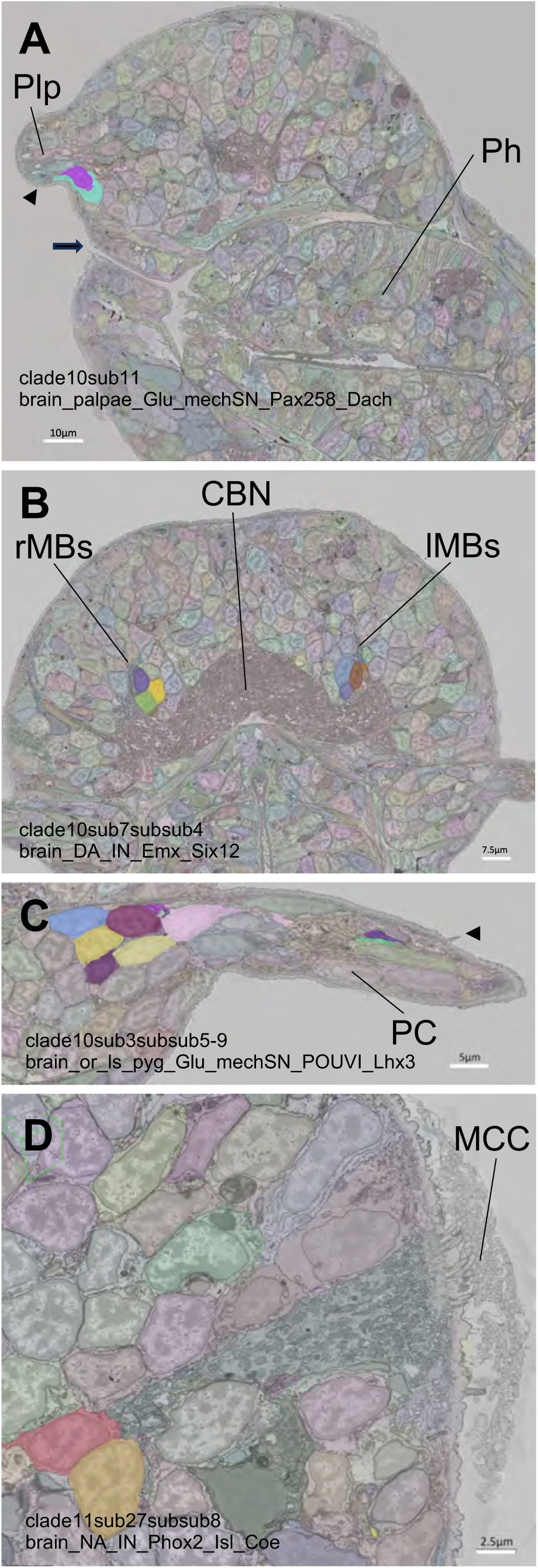
Mechanosensory cell types. **A.** Localisation of Clade10sub11 to the palpae. Plp, palpae; Ph, pharynx; arrowhead, position of mechanosensory cilia; arrow, mouth opening. **B.** Localisation of Clade10sub7subsub4 to the base of the mushroom bodies. rMBs, right mushroom body stalk; lMBs, left mushroom body stalk, CBN, central brain neuropil. **C.** Localisation of Clade10sub35subsub5-9 to the base of the peristomial cirri (PC). Arrowhead points at mechanosensory cilium. **D.** Localisation of Clade11sub27subsub8 in juxtaposition to multiciliated cell (MCC).

Motile mechanosensory cilia are also present in three neuron types of the *FEZF+* family, two of which (clade10sub9 and clade10sub15) specifically express Pkhd1l1. *FEZF+* neurons are located at the base of the antennae and of the parapodial cirri. One *FEZF+* neuron type (clade10sub9) expresses an extra duplicate of TrpV; another (clade11sub45subsub0) expresses *r-opsin1* and possesses a long sensory process, which we identify as a rhabdomeric photoreceptor in the position of the late larval eye. The closely related *Goosecoid/Fer2* family shares with the *FEZF\** family members expression of the Nkx2.1 transcription factor (Fig. 4A). Significantly, members of this family express markers for catecholamine and histamine synthesis and transport, highly reminiscent of the dopaminergic *Fer2+* neurons in nematodes and flies ^56^. For example, neurons located at the apex of the brain express enzymes for noradrenalin and tyramine production and are loaded with electron-dense secretory vesicles.

A third family of mechanosensory cell types expresses *PouIV*. Among these, one cell type (clade10sub3sub5-9) uniquely expresses *PKD1.1* and *PKD2.1*, encoding the PKD1/PKD2 heteromeric polycystin complex, and BAIAP, which is known to play a role in the regulation of stereocilia in hair cells. We have identified these as glutamatergic ciliated sensory neurons that are located at the base of the antenna, peristomial cirri, and anal cirri (Fig. 6C). Another glutamatergic *POUIV+* cell type (clade10sub21) with sensory cilia and a unique TrpV2 homolog is also found in the peripheral ganglia. These *PouIV+* neurons correspond to the “collar receptors” of the “mechanosensory girdle” ^57^. We find the *PouIV+* mechanosensory neurons closely related to another family of neurons, the *Phox2+* neurons, which are located adjacent to the *PouIV+* neurons in the peripheral ganglia of the larval segment and co-express the HAND bHLH transcription factor (e.g., clade11sub22subsub4). In contrast to the *PouIV+* neurons, the *Phox2+* neurons are cholinergic and do not form sensory cilia. Importantly, the only neuron type with a clear noradrenergic profile (co-expression of tyrosine hydroxylase and dopamine ß-hydroxylase) forms part of the *Phox2+* family. These neurons are consistently found at the base of multiciliated cells in proto- and telotroch and/or of muscle cells, consistent with a proprioceptive function (e.g., Clade11sub27subsub8; Fig. 6D).

Another prominent family of brain neurons shows expression of the transcription factors *Dlx* and *KLF4*, together with *Six4, rx*, and *Bsx*. These represent a family of previously characterised sensory-neurosecretory cells known as “apical nervous system” (ANS)^58,59^. Here, we find that ANS neuron types specifically and consistently express a variety of opsins known to play a role in ambient light detection, including OPN3 (encephalopsin), OPN5 (neuropsin), RRH (peropsin), and Go-opsin orthologs (Supplementary. Figure 9). This goes in concert with the specific expression of numerous vertebrate-type olfactory receptors (ORs), of which we find a total of 33 putative paralogs indicating independent massive duplication of these receptors in the annelid (Supplementary Figure 10. Focussing on *Bsx+* neuron types, these assemble into lateral sensory organ-like structures with chemosensory non-motile cilia (validate) (FIg. 6g). Intriguingly, the massive expression of different opsins also extends to another neuronal family characterized by the expression of *Rfx4*, a gene supporting primary cilia formation^60^.

## Discussion

### A whole-body single cell object mapped to an EM volume

We present the first resource mapping all differentiated cell types of an animal to an EM volume. This opens up new avenues towards understanding how cell type-specific gene expression translates into cellular features including molecular machinery, physiology and shapes. Genes encode specific receptor and response modules, specific pathways, cytoskeletal elements, endo- or lysosomes, excretory vesicles, or entire organelles that characterize cell types. Our resource enables an organism-wide understanding of this process, and thus addresses the cellular genotype-phenotype link in unprecedented resolution.

Previous attempts to link gene expression to EM volumes focused on the correlation between spatial transcriptomics and EM in specific brain regions, to map transcriptional and ultrastructural responses^61^. Serial lift-out applied to *C. elegans* larvae yields a cryo-ET dataset sampling the worm’s anterior-posterior axis, and thus enables the study of multicellular molecular anatomy^62^. The largest EM volumes to date were produced for connectomics^63^. Although these invaluable datasets have been combined with neural activity^64^ and, in part, transcriptomic data^65^, integration has generally focused on the extracted connectome, disregarding ultrastructural detail. Our resource and integration methods, by contrast, allow molecular information to be directly embedded within the EM volume’s structural framework.

### Large coregulons define major cellular phenotypes

We have devised an approach to systematically screen for genes co-regulated across our curated differentiated cell types by positioning genes in cell type space, where expression in a given cell type represents one dimension. Our data reveal cluster of genes coregulated across few to many cell types, which jointly implement characteristic cellular features. Similar gene sets are also obtained from weighted gene co-expression network analysis (WGCNA). From these analyses, we uncover 10 large coregulons that encode major cellular phenotypes including stem cells and germ line, epidermal cells, digestive cells of the midgut, mesothelial cells of the cardiovascular system, myofibres characteristic of somatic musculature, glia, glands or neurons. Others encode one large organelle such as the motile cilium that is either present or absent. Coregulons appear crucial for tissue formation and thus for the overall organisation of the body. The large coregulons include few highly conserved transcription factors, which are active in all cell types that activate the coregulon, and we show that binding sites for these transcription factors are enriched in the genes of the coregulon, indicating direct regulation for at least many of these genes. Many of these factors are known to be involved in the specification of similar tissues in vertebrates or insects, such as grainyhead for epidermis^66^, HNF4 for digestive tis^67^; MyoD for muscle^68^, FoxF1and serum response factor for smooth muscle^69^, Glial Cell Missing (GCM) ^70^ and NR4A^71^ for glia, FERD3L for glands^72^, Rfx and Foxj for multiciliated cells^73^, and Myt1L for neurons^42^. This makes it highly likely that these coregulons and the tissues they specify are conserved at least across bilaterians and were present in Urbilateria.

At least some of the coregulons in *Platynereis* are not mutually exclusive, but can be co-activated in one and the same cell type. In particular, this is true for the motile cilia coregulon and the digestive coregulon, which can co-occur in neurons. The neuronal coregulon is also special in that it appears to differ from other coregulons in its regulation. While other coregulons express conserved transcription factors known to activate gene expression, in line with the presence of their binding sites in other genes of the same coregulon, we could only identify one truly pan-neuronal transcription factor, Myt1l, which is known to act as a repressor of non-neuronal fates in other systems^42^. None of the other transcription factors that form part of the neuronal coregulon is expressed in all neurons; the most widespread factors being LIM and POU as well as the Cux/onecut honeodomain factors ^74^.

### Evolutionary conservation of neuronal families?

Focusing on neuron types, our study reveals that these are defined by the differential expression of homeodomain and bHLH transcription factors, in line with other studies^75^. Our systematic screening for highly differentially expressed transcription factors has revealed cell type-specific combinations of transcription factors that define neuronal types. Similarities in transcription factor expression across types have then allowed us to define 14 neuronal families that express family-specific combinations of transcription factors (Figure 4; Supplementary Figure 4), and the combination of cellular-resolution transcriptomics and volume EM available in the new *Platy*Browser enabled the molecular and morphological characterization of representative neuron types within these families. These results are especially relevant from a comparative angle, because the 6dpf neuronal families in *Platynereis* match neuronal types and families that have been identified previously by developmental neurogenetic studies as possibly conserved between vertebrates, insects and nematodes.

First, we find *Mnx+* cholinergic motor neurons, as well as *Dbx+* GABAergic inhibitory and *Evx+* excitatory commissural interneurons, as main constituents of the *Platynereis* CNS. *Mnx+* motoneurons exist in insects and vertebrates^44^ and were also found in *Platynereis*^40,76^ and in molluscs^77^. They were most likely cholinergic in Urbilateria, with Isl1-Lhx3 complexes activating the key transcription factor mnx^78^. In addition to segmental *mnx+* neurons, we find a Mauthner or VeLD-neuron-like^79^ *mnx+* brain motor neuron type with contralateral descending projections along the longitudinal muscles, which corresponds to the brain motor neurons described for the 3dpf *Platynereis* young worm that mediate body bending in response to phototaxis^80^. This would suggest that one ancient function of the *Mnx+* motor neuron family may lie in body bending. Inhibitory *Dbx+* GABAergic commissural interneurons are known as V0d interneurons in vertebrates and control left-right or agonist-antagonist alternation of muscular contraction via direct inhibitory input to motoneurons^81^. In the fly, local *Dbx+* interneurons coordinate flying and walking^82^. *Dbx+* neurons were previously described for *Platynereis*^40,76^ and have now been identified as GABAergic and commissural. *Evx+* glutamatergic neurons are likewise known for vertebrates as the V0v interneurons^83^. Acting downstream of the *POUIV+* Rohon-Beard neurons, they trigger contraction contralateral to the site of mechanical stimulation. In *Drosophila, Evx+* commissural interneurons coordinate bilateral locomotor patterns^84,85^, and we now find that commissural glutamatergic *Evx+* neurons also exist in the annelid. Intriguingly, in vertebrates the *Mnx+*, *Dbx+*, and *Evx+* neurons are the main constituents of the trunk central pattern generator (CPG). We take the co-conservation of these families in the *Platyereis* 6dpf young worm as evidence that a such-composed CPG may have existed in the Urbilateria.

Second, we find that in *Platynereis* the *Dbx+* commissural interneurons belong to a larger family of GABAergic interneurons (box in Fig. 4) that likewise appears evolutionarily conserved. Neuron types of this larger family have in common to express the bHLH factor *Ptx1a* together with the CSL family member Rbpj, and the exact same combination holds true for vertebrate *Ptx1a+* GABAergic inhibitory neurons that are present along the length of the neural tube, in the retina^86^, the arcuate nucleus of the lateral hypothalamic area^8^, the cerebellum^9^, the trigeminal nuclei of the hindbrain^10^, and in the spinal cord^11^. Remarkably, in *Platynereis*, *Dbx, Ptf1a,* and *Rbpj* are co-expressed (while the two families are strictly separate in the vertebrates), which suggests that GABAergic interneurons may have initially evolved as one family with *Dbx+, Ptf1a, Rbpj* identity. Strikingly, in *Platynereis* the two mushroom body neuron types from part of this family, too, indicating that the annelid sensory associative centers evolved as a brain specialization of the ancient *Dbx+, Ptf1a, Rbpj+* inhibitory GABAergic interneuron family.

Third, our in-depth molecular and morphological analysis of *Platynereis Pou4+* neuron types^40^ reveals that these are mechanosensory with motile cilia and microvilli very similar to hair cells, which are considered evolutionarily ancient^40,77,87^. In vertebrates, *Pou4+* mechanosensory neurons are represented by the *Rohon-Beard* primary sensory neurons in basal vertebrates^88^, and by the functionally redundant neurons of the dorsal root ganglia^79^. *POU4+* mechanosensory neurons also exist in mollusks^77^ and nematodes^89^. Furthermore, in our analysis the *Pou4*+ mechanosensory neurons are most closely related to the *Phox2+* proprioceptive neurons (box in Figure 4x). In both annelids and vertebrates, these neurons are cholinergic and noradrenergic, and their direct vicinity to multiciliated cells and myofibers in the 6dpf suggests direct sensory and modulatory interaction with these cells. This indicates evolutionary conservation of the *pou4+* somatosensory versus *phox2+* viscerosensory dualism as previously suggested ^77,90^.

### Serially homologous neuron types in a circular nervous system

Strikingly, in the Platynereis nervous system all possibly ancient neuronal families have in common that individual neuron types that belong to these families are found from anterior to posterior in all six segments of the body (Fig. 4b, c). For example, the *Mnx+* motor neurons*, Dbx*+ and *Ptf1a+* GABAergic commissural interneuron, and *Pou4+* mechanosensory neuron types discussed above are all present in brain and nerve cord, and the same holds for most other families. This observation corroborates the notion that bilaterian neuron types first evolved in a concentric arrangement as part of a circular nervous system, and were then arranged along the evolving bilateral nerve cords and brain in early bilaterians, so that they now occur at all body levels. Their subsequent specialization in the trunk towards locomotor and in the head towards behavioural control functions has shaped the centralisation of the nervous system towards nerve cords and brains. In Fig. 7, we present a first hypothesis on how nervous system centralisation might have taken place in early bilaterians, taking into account the serial arrangement of homologous neuron types in a ring around the midline of the body.

**Figure 7.**
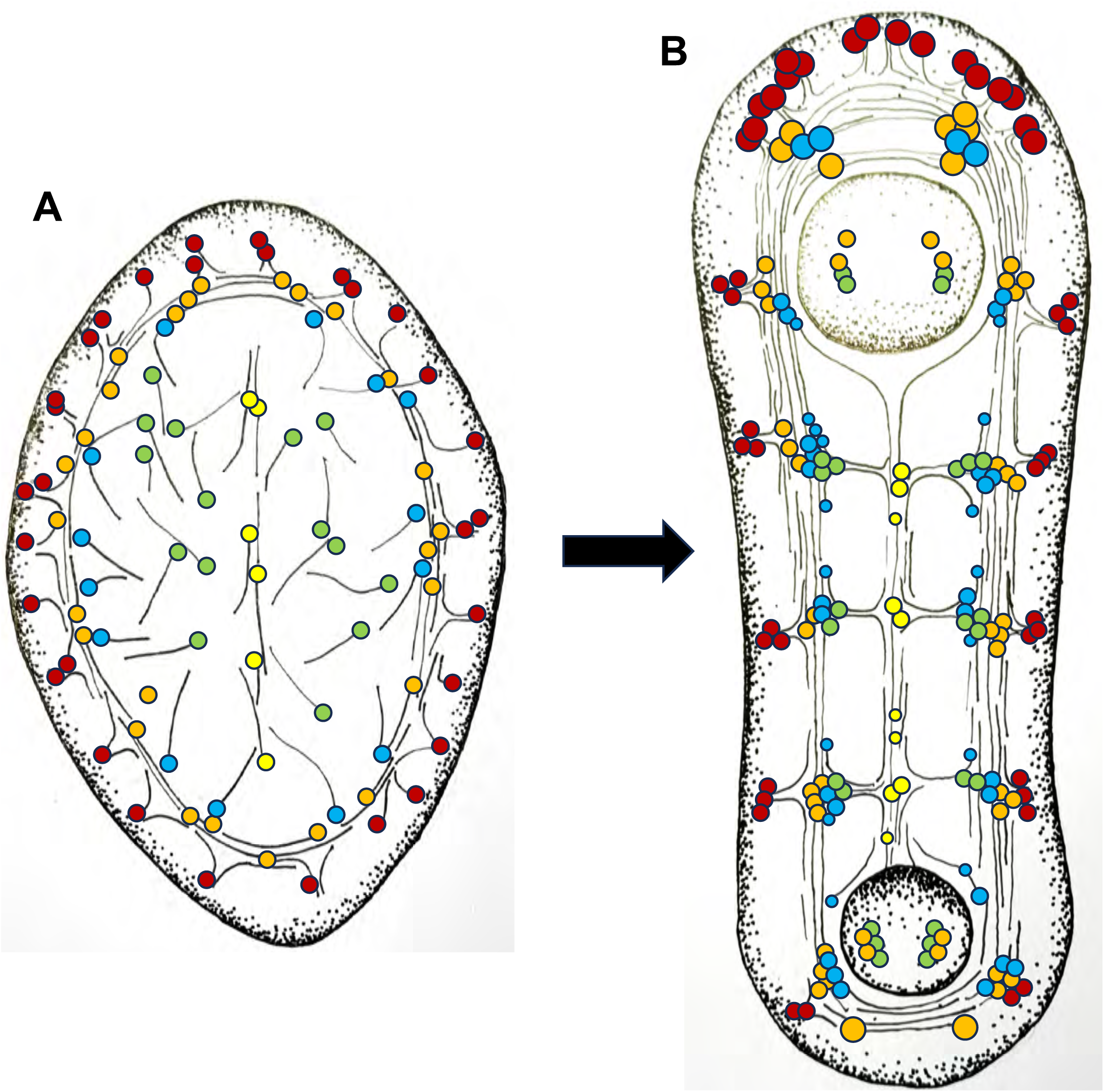
A new hypothesis for nervous system centralisation. **A.** Non-centralized nerve net in a hypothetical animal in the bilaterian stem. Ventral view with cholinergic (green) and serotonergic (yellow) motor neurons being part of a mucociliary feeding sole as was present in *Dickinsonia*. Mechanosensory neuron types (red) are located towards the periphery of the animal. Neuron colours indicate gene families from Figure 4. **B.** First CNS in Urbilateria. Neurons of the *Dbx, Ptf1a+* (blue) and *Evx+* (orange) interneurons families together with the *Mnx+* (green) motor neurons have aggregated as segmental central pattern generators (CPGs). Mechanosensory neurons (red) in the periphery form peripheral ganglia. Neurons in the first segment of the body aggregate as brain rudiment including first sensory associative centers, Note that motor and interneurons of the oral and most posterior body segment move inwards as enteric nervous system with the internalising anterior and posterior gut rudiment.

This principle of serially homologous neuron types in a compressed ring-shaped arrangement has long been recognized in the vertebrates and underlies major attempts to systematically understand nervous system architecture, exemplified in the comparative work of Swanson^91^ and in the prosomeric model of Puelles and Rubenstein^92^. In essence, these authors identify components of the basal plate, which harbours motor components, and of the alar plate, which harbours sensory components, along the length of the neural tube. The serial homology and the lateral-to-medial arrangement of sensory and motor components, deduced from a nerve net-like arrangement, is central to our model, which may thus serve as a first stepping stone towards understanding nervous system centralisation.

## Supporting information

Combined supplementary data

## Supplementary Figures

**Supplementary Figure 1:**

A) UMAP embedding of all nuclei of the atlas is colorcoded by clade signature.

B) Canonical marker gene expression across the clades is visualized as a violin plot.

**Supplementary Figure 2:**

List of HCR registered genes in the new version of PlatyBrowser with XLOC ID and probe transcript for reproducibility.

**Supplementary Figure 3:**

**A)** Experimental strategy to find clusters of co-expressed genes.

**B)** tSNE embedding of genes with Louvain clusters indicated by color.

**C)** Genes grouped to clusters were used to score cells in the differentiated atlas. Selected gene cluster scores are presented as features on the atlas UMAP.

**D)** Hierarchical clustered dotplot visualizes all averaged scores across the clades.

**E)** Averaged scores across terminally differentiated cell types are shown. Order of celltypes follows the Neighbor joining tree in Fig 1.

**F)** Genes used for the WGCNA analysis are indicated on the tSNE embedding and are grouped by the WGCNA modules they belong to.

**G)** Venn diagram visualisation of selected gene clusters against WGCNA gene modules shows substantial overlap between the gene groups identified by independent methods.

**Supplementary Figure 4:**

**A)** Expression pattern similarity between coregulons

**B)** Correlation between coregulons and cell types

**C)** Expression of POU, LIM, and Cux/Onecut homeodomain, and of the Myt1L repressor against all curated cell types. Note widespread expression in neuronal cell types.

**Supplementary Figure 5:**

Larger version of Figure 4A with cell type and gene names.

**Supplementary Figure 6:**

Synthesizing enzymes and release channels for individual transmitter systems as detected in the *Platynereis* genome.

**Supplementary Figure 7:**

Expression of enzymes for transmitter synthesis and channels for vesicle release plotted against curated neuronal cell types

**Supplementary Figure 8:**

Expression of mechanosensory and warmth-sensitive Trp channels plotted against curated neuronal cell types

**Supplementary Figure 9:**

Expression of opsins plotted against curated neuronal cell types.

**Supplementary Figure 10:**

Expression of chemosensory and olfactory GPCRs plotted against curated neuronal cell types.

**Supplementary Figure 11:**

Expression of neuropeptides plotted against curated neuronal cell types

## Acknowledgments

We thank Christian Tischer of the Bioimage Analysis Support Team at the European Molecular Biology Laboratory (EMBL) for support. Christian Tischer has been supported by grant number 2020-225265 from the Chan Zuckerberg Initiative DAF, an advised fund of Silicon Valley Community Foundation. We thank the Advanced Light Microscopy Facility (ALMF) and Genecore at the European Molecular Biology Laboratory (EMBL) and Olympus and Leica for support.

The work was supported by the European Research Council Advanced grant 788921/NeuralCellTypeEvo to D.A. and by a data creation award from the Life Science Alliance. CDC is supported by an European Molecular Biology Organization (EMBO) Postdoctoral Fellowship (ALTF 186-2023). A.B.K. receives funding from Horizon Europe Marie Curie Doctoral network ZooCELL under grant agreement no. 101119891.

## Methods

### Nuclei Isolation

Nuclei were extracted based on the protocol described by the Kaessman laboratory^93^. 6dpf Platynereis larvae were immobilised with NoCa-NoMg ASW and collected in 1.5ml LoBind tubes. For snRNAseq, the larvae were flash frozen in liquid nitrogen and stored at −80°C before use. For bulk ATACseq, nuclei from the larvae were extracted directly afterwards. Frozen or fresh larvae were then placed on ice and homogenized in 300 µl of HB buffer (250 mM sucrose, 25 mM KCl, 5 mM MgCl₂, 10 mM Tris-HCl pH 8.0, 0.1% IGEPAL, 1 µM DTT, 0.4 U/µl Murine RNase Inhibitor [NEB], 0.2 U/µl Superase-In [ThermoFisher] and 1x eComplete Protease Inhibitor (Roche) using micro pestles (Axygen). Homogenization was performed on ice by hand and completed within 5 minutes. Unlysed tissue fragments were pelleted by centrifugation at 100 × g for 1 minute at 4 °C. The supernatant containing nuclei was transferred to a new LoBind tube and centrifuged at 400 × g for 4 minutes at 4 °C. After discarding the supernatant, the pellet was washed and resuspended in 300 µl of fresh HB buffer and nuclei were pelleted again at 400 × g for 4 minutes at 4 °C. Finally, the nuclei pellet was resuspended in 50 to 100 µl of ice-cold Dulbecco’s PBS supplemented with 1 µM DTT, 0.4 U/µl Murine RNase Inhibitor (NEB), 0.2 U/µl Superase-In (ThermoFisher) and 1x eComplete Protease Inhibitor (Roche). The suspension was passed through a 40 µm Flowmi filter to remove aggregates. Nuclei quality and concentration were assessed by staining with SYBR Safe DNA stain and visualisation on a Zeiss Axio Imager using a C-Chip hemocytometer (Neubauer improved).

### Single Nuclei Capture and Library Preparation

Single nuclei capture and library preparation were performed using the Chromium Single Cell 3’ Gene Expression Kit (v3.1 chemistry) and the Chromium Controller (10x Genomics), following the manufacturer’s protocol with slight modifications. Approximately 15,000 nuclei were loaded per capture channel. cDNA amplification was performed with 15 cycles for samples at stage 29–30 and 12 cycles for samples at stage 30–31, using a 3-minute extension time. Final libraries were quantified using a Qubit fluorometer, and fragment sizes were assessed using an Agilent Bioanalyzer. Libraries were sequenced with a minimum of 10,000 PE reads per nuclei using the following read configuration: 28 cycles for Read 1, 10 cycles each for the i5 and i7 indices, and 90 cycles for Read 2.

### Single Nuclei Data Processing and atlas generation

The single-cell atlas was generated from six larval batches originating from independent matings (2–4 technical replicates), yielding 14 cDNA libra fastq files were mapped against the *Platynereis Dumerilii* genome 2.1^94^ using CellRanger v.6 (10X genomics). Only nuclei considered as cells by CellRanger were used for the atlas generation and cells needed to have at least 200 genes detected. Additionally, doublets were filtered using Scrublet^95^. All snRNA-seq data processing was performed using Seurat v4^96^. The data was scaled across all genes with regressing out effects of differential RNA counts and detected mitochondrial genes and 200 PCs were calculated in a subsequent PCA analysis. To remove batch effects, the different libraries of the atlas were integrated with Harmony using 150 input PCs^97^. Clustering using default parameters was performed with the Louvain algorithm using 150 Harmony components. To group clusters in higher order classes, we constructed an unrooted neighbor-joining tree (APE package in R) from averaged transcriptomes of the initial clusters. Thirteen clades with >80% bootstrap support were identified; clusters outside well-supported clades were grouped as “no clade” for manual inspection.

Each clade including a no-clade group was re-normalized with SCTransform in Seurat, re-integrated using Seurat’s anchor-based method, and clustered with Louvain, respectively. For subsets with too few cells we lowered the subclustering parameters, and one very small clade (clade8, 113 cells) was not further subclustered. Resulting subclusters were manually inspected for following criteria. i) The top 120 markers were evaluated for transcription factor expression, ii) their specificity relative to related populations (“individuation”), and iii) their overall marker gene plausibility based on literature. Clusters not fulfilling these criteria were removed from the data set. Neuronal clades underwent additional refinement, including merging of over-split populations and further sub-subclustering when additional heterogeneity remained such as Tal-Gata cells^40^. After completion of clade-based subclustering and curation, all validated populations were reassembled into a single atlas. This curated dataset represents the fraction of cells with clear terminal differentiation and interpretable transcriptomes, suitable for mapping to the Platybrowser resource

### Bulk ATACseq and Library Preparation

Chromatin accessibility was profiled using an OmniATA optimized for platynereis^98^. 50000 nuclei were permeabilised with ATAC-RSB/NTwD for 3 minutes before washing with ATAC-RSB/Tw. Nuclei were then pelleted and resuspended in a 50 µL of transposition mixture containing (100nM loaded Tn5, 0.01% digitonin, 0.1% Tween-20, 10mM Tris-HCl, 10mM MgCl_2_, 10% DMF). Tagmentation was carried out at 37 °C for 30 min with gentle agitation and stopped by addition of 5 volumes of DNA binding buffer (Zymo). DNA was purified using Zymo Clean and Concentrator-5 kit and eluted 25ul EB buffer. PCR was done with 2x NEBNext HIFI master mix and Nextera i5 and i7 primers for dual indexing and amplified for 10 cycles. Libraries were quantified and checked by Qubit and D1000 Tapestation. Sequencing was done with 75PE on Nextseq2000 for 50million reads.

### Read processing and peak calling

Raw ATAC-seq reads were processed using the OpenChromatin pipeline (Zaugg Lab, EMBL; https://github.com/zaugglab/openchromatin-pipeline), which provides a standardized framework for read alignment, QC, and peak calling. Reads were checked with FastQC and adapter-trimmed using Trim Galore!. Trimmed reads were then aligned to the platynereis v2.1 genome using Bowtie2 with parameters optimized for short ATAC-seq fragments. Duplicate reads and mitochondrial alignments were removed with Picard and Samtools, and fragment length distributions were computed for QC evaluation. Peak calling was performed using MACS2, and replicate reproducibility was assessed via IDR analysis to produce a consensus set of high-confidence open chromatin peaks.

### Coregulon motif enrichment and discovery

Peaks were assigned to the nearest annotated genes and aggregated into coregulon modules. Motif discovery was performed on coregulon-associated peak sets using the MEME-Suite (v5.5.5). Over-represented motifs were identified with AME against the JASPAR 2024 CORE and Metazoa motif databases. Both “global” and “pairwise” enrichment analyses were performed using Fisher’s exact test with average odds scoring. Significance values (−log_10_ *p*) were extracted from the AME text outputs and aggregated across modules to generate a matrix of motif enrichment values. Motif discovery was carried out on repeat-filtered sequences within 200 bp windows centered on peak summits. The resulting motifs were compared against the JASPAR 2020 CORE Metazoa database using TOMTOM to assign putative transcription factor family identities based on similarity of position weight matrices.

### HCR Staining, sample preparation and imaging

*Platynereis dumerilli* were collected at exactly 6 days post fertilization (dpf) in ASW, immobilized with NoCa-NoMg ASW, and then incubated with Proteinase K for 3 min in order to digest the cuticle. The larvae are then fixed in 4% PFA for 30 min, washed in PTW, and dehydrated in stepwise 25% increasing concentrations of EtOH for long-term storage at −20°C. After stepwise rehydration in PTW from storage, larvae were stained with probes purchased from Molecular Instruments Inc. following an established whole-mount multiplexed embryo protocol^99^. After successful HCR staining, the larvae were incubated with a 1ug/ml DAPI concentration for 30 minutes to stain nuclei for registration. Upon completion of the full staining protocol, larvae are embedded in increasing volume of SlowFadeGlass Antifade mounting medium in (at least) 15 min steps to avoid rapid deformation of individuals due to the insertion into a hydrophobic environment, and are left in this mounting medium overnight for acclimatization purposes.

Embedded larvae are then mounted on a special slide construction for imaging. 0.12mm spacers are placed between two #1.5H coverslips for two-sided imaging, and are then taped to a regular glass microscopy slide. Imaging was performed on Leica SP8, Stellaris 8 (WLL) and Olympus iXplore SPIN SR confocal and spinning disk confocal microscopes. All imaging was done on 40X Oil Plan-Apo objectives (on both Leica and Olympus microscopes). Full-body volumes of larvae were acquired in 0.5um Z-steps in 3 channels covering DAPI, Alexa-546 and Alexa-647 fluorophores.

### Light microscopy to EM volume registration

The registration pipeline takes as an input the PlatyBrowser SBEM volume nuclei segmentation, as well as two-sided (dorsal and ventral) confocal imaging 3D volumes in DAPI and gene channels. Only the DAPI channel was used for registration, with the final transformation being applied to the gene channels. The two opposing confocal views were combined using a three-step registration procedure consisting of pre-alignment, rigid alignment, and deformable alignment, culminating in a final composite volume. Pre-alignment was necessary due to the random orientation of animals during imaging. Both images were smoothed using thresholds to obtain a point cloud. Then PCA was applied to the point cloud and the image was rotated to align the first principle component with the X axis. After that a heuristic approach was used to ensure that all animals have the same orientation, with the head of the animal oriented towards 0 due to the fact that the head of the animal has more signal than the rest of the body. Pre-aligned images were registered using Euler transform in Elastix. Firstly, the registration was done using a mutual information metric with 5 levels in the image pyramid schedule (64, 32, 8, 4 and 1) and 20000 spatial samples. In the second step rigid registration was done using normalized correlation coefficient using 5 levels in the image pyramid schedule (16, 8, 4, 2 and 1) and 50000 spatial samples for metric and gradient estimation. Due to small deformations resulting from the sample handling between the acquisition of the views it was not possible to fully register two views using only rigid registration. Elastix B-Spline deformable registration was used as the last registration step. The metric consisted of normalized correlation, mutual information and rigidity penalty with the corresponding weights of 1, 1 and 100. Registration was performed in one step at the original resolution with the final grid spacing of 64 voxels in X and Y directions and 16 voxels in Z direction using 100000 spatial samples for metric and gradient estimation. Small inaccuracies in the registration of two views can cause duplication of the spots, and thus instead of directly averaging the registered views, a composite volume was constructed. The DAPI channel of both registered images was smoothed with gaussian with kernel size of (10, 20, 20) (ZYX). Then the volume was split into the areas where one smoothed view had a higher intensity than the other and this mask was used to create a weighted average of the views.

Each composite volume was segmented with Stardist and mutex watershed. Thanks to the strong shape prior, Stardist could detect separate nuclei in the areas of the sample where resolution in Z did not allow to reliably predict a boundary between separate nuclei. Mutex watershed was able to accurately segment nuclei of unusual shape and generally segmented the foreground more precisely. The instance segmentations obtained with these two methods were combined resulting in ∼ 10500 nuclei detected in each sample. Despite the errors, especially caused by autofluorescence, the segmentation was consistent throughout the volume which is important in order to use it as a source of registration landmarks. Spot detection was performed using a custom 3D Spotiflow model^100^. Having a robust model for spot detection allowed for the quantification of gene expression signal and straightforward aggregation of the signal from different replicates despite intensity variations across and within the samples. The volume was split into regions by the distance to the closest nucleus, approximating the cytoplasm of the cell. After that the number of spots detected in each region was calculated and assigned as the gene expression value to each nucleus.

As a next step, all confocal volumes were registered to the EM volume, which was used as a template. To achieve this, EM segmentation and corresponding centroid point clouds were set as fixed. As a result, the displacement field and nucleus-to-nucleus matching between each nucleus in each confocal volume and the nuclei in the EM volume were outputted.

This part of the pipeline is fairly generic and can be applied to different biological samples, given that the instance segmentation is already performed.

Before doing a final nucleus-to-nucleus deformable alignment, it is necessary to rigidly align the samples to the template EM volume as closely as possible. In this case similarity alignment (translation + rotation + scaling) was used to compensate for the size difference caused by different fixation and staining protocols. After this step of the pipeline, samples are registered as closely as possible without deformable alignment. Both EM and light segmentations were then converted to point clouds by extracting coordinates of centroids. Coherent Point Drift^101^ was used to perform deformable alignment with the EM point cloud as a fixed set and light microscopy point clouds as moving point sets, resulting in the aligned point clouds. Coherent Point Drift finds the coordinates of points of the aligned point cloud, so an additional matching step was implemented to find a correspondence between light microscopy and EM nuclei. The total distance between matched nuclei was minimized using CVXPY with a constraint that each nucleus in both datasets should match one or more nuclei in the other dataset. This matching enabled annotation transfer between the datasets, including for example, assignment of gene expression based on spot detection to nuclei in EM.

The final result of registration is the matching between nuclei in confocal volumes and the EM template. It is very hard to assess the registration quality based on the point clouds, therefore a final phase was added to the pipeline to overlay raw fluorescence data with the electron microscopy. Deformable transformations were found using Elastix and applied to the nuclei segmentation as well as all channels of the raw confocal volumes. This deformable registration with matched nuclei centroids as landmarks was then used to warp the confocal images and overlay them with the EM data.

Registered instance segmentation as well as all channels of the raw confocal volumes and the final gene expression table were added to the existing MoBIE project. This allows users to interactively visualize multiple datasets together, perform arbitrary direction slicing and visualize the mapping of the quantified gene expression signal to the segmentation masks.

### Automated annotation of cell types to EM

The automatic annotation pipeline can be split into four separate parts: preprocessing, profile creation, probability assignment, and visualization. Preprocessing involves cleaning the imaging (ProSPr and HCR registration) data in order to remove unnecessary complexity for computation, and then binarization to generate final gene expression tables per nucleus.

snRNAseq preprocessing also involves subsetting and data curation for the creation of pseudobulk gene expression profiles of cell type clusters that are then used in the probabilistic assignment. Profiles “zero” and “autofluorescence” are also created to make the assignment of extremely low-expressing or high-expressing cells more robust. Probabilistic assignment was then calculated in the following manner:

Given: a scRNAseq atlas consisting of 𝐺 genes and 𝑁 cells split into 𝑁_𝑐𝑒𝑙𝑙𝑠_ clusters, and for gene 𝑘 number of cells in which this gene is expressed in cluster 𝑖 is 𝑛𝑜𝑛𝑧𝑒𝑟𝑜_𝑘,𝑖_ ∈ [0; 1]. With input of: cell in EM with expression vector 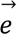, for example, (0, 0, 1, 1, 0), so binary measurements “expressed / not expressed” for a subset of genes of the original atlas, 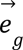 is the binary number corresponding to the gene 𝑔. Gives output: probability that given the expression measurements the cell belongs to cluster *i* ^P^ ^(cell^ ^∈^ ^cluster^ ^i^ ^|^ ^e^ ^→), i^ ^∈^: 1 … N

Method: Using Bayes theorem

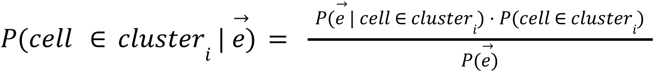

All parts can easily be estimated from the scRNAseq atlas:

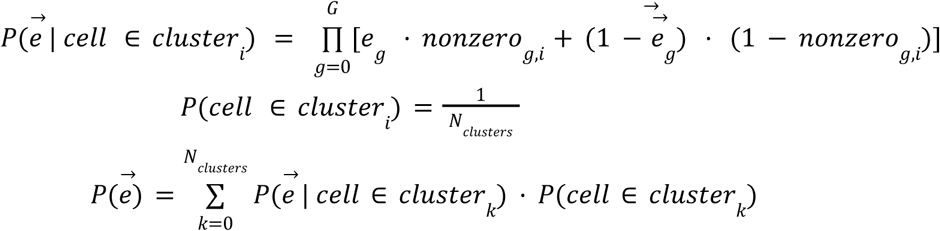

Resulting in a table of size 𝑁_𝑐𝑒𝑙𝑙𝑠_ × 𝑁_𝑐𝑙𝑢𝑠𝑡𝑒𝑟𝑠_ where each element is a probability that the given cell belongs to the given cluster.

The impact of any gene can also optionally be weighted by specificity for the purpose of increasing the importance of very lowly expressed but specific marker genes. To assess the results of the assignment (Fig.2D), a scaling for visualization step is provided where the most probable clusters for any given cell are ranked.

### Cloning and detection of the *mnx*-repoter construct expression

4181 bp of the regulatory sequence upstream of the translation start site of mnx were cloned into the Palm-3xHA-tdTomato P2::H2B-GFP^102^ plasmid using SalI and SgsI restriction sites to generate the mnx-reporter construct. Transgene construct was confirmed by Sanger sequencing. 1 or 2 cell embryos were injected with the empty plasmid (control) or with the *mnx* reporter construct following standard procedure. GFP expression was used to screen for positive larvae at 48hpf. Larvae were stained using anti-hemagglutinin antibody (rabbit-Abcam, dil 1:1000) with rhodamine-phalloidin and imaged with Leica TCS SP8 microscope using a 63x objective.

### CRISPR KO

#### gRNA design

gRNAs were designed using online tools. The online tool “ZiFit” (http://zif.partners.org) was used to design the dbx gRNA (AACTCGGTCCTTGCCTC<u>TCC</u>). “sgRNA scorer 2.0”^103^ was used to design the neogenin gRNA (GATGGCCGCTGCAAGGTCCC<u>TGG</u>). gRNA sequences were blasted against platynereis genome and transcriptome to avoid SNP region and ensure specificity. The gRNA sequences were ordered as ssDNA and annealed using a slow cool-down approach from 95◦C to room temperature in annealing buffer (3µl of 100µM oligos in 44µl of buffer). The resulting dsDNA were cloned into the vector pDR274 (Addgene -Plasmid 42250), linearized with BsaI. gRNA were generated from in vitro transcription of DraI-linearized construct using the MAXIscript Kit (Thermo).

#### Embryo injection

Knock out (KO): The gRNAs (Dbx gRNA or Neo gRNA) were microinjected into the 1-2 cell larvae together with mRNA coding for the Cas9 protein. Dextran fluorophore was added to the injection cocktail in order to screen for successfully injected larvae. In the dbx experiment, no-gRNA control was used. For the neo/dcc experiments, a scrambled sgRNA was used as control. Only larvae with appropriate development with strong fluorescent dextran signals were selected for analysis.

### Analysis of mutants

Standard nested PCRs were performed from genomic DNA using two pairs of specific primers.

Neo/DCC genotyping oligo 1 (5’-3’) AGTTTCTGTGGACTCCTTCGA

Neo/DCC genotyping oligo 2 (5’-3’) CGGCTTTGGTCTTCACAGAG

For fast genotyping the dbx mutants, the nested PCR products were purified and digested using AvalI, as the gRNA was designed to target a region containing a unique restriction enzyme site. Digested product was purified and size differences were analyzed on a 3% MetaPhor Agarose gel.

For both dbx and neo candidate mutants, the nested PCR products were cloned into the Topo TA vector, and individual clones were sent for sanger sequencing. A great variety of insertions and deletions were detected in the case of dbx. For neogening a small deletion was detected.

### Commissural axon quantification

Larvae were grown at 18°C and fixed at 50hpf, when the ventral nerve cords are furthest apart and the commissural axons most visible. Animals were then immunostained with an antibody against acetylated tubulin to visualize the commissural neurons. Larvae were imaged ventrally on an Sp8 confocal microscope. Maximum projections were produced from the obtained z-stacks and the axons in the VNC were manually counted. Comparisons were tested using ANOVA test followed by Tukey’s1.

