## Supplementary material for "A whole-body cell type atlas mapped into an electron microscopy volume of an annelid worm": Combined supplementary data

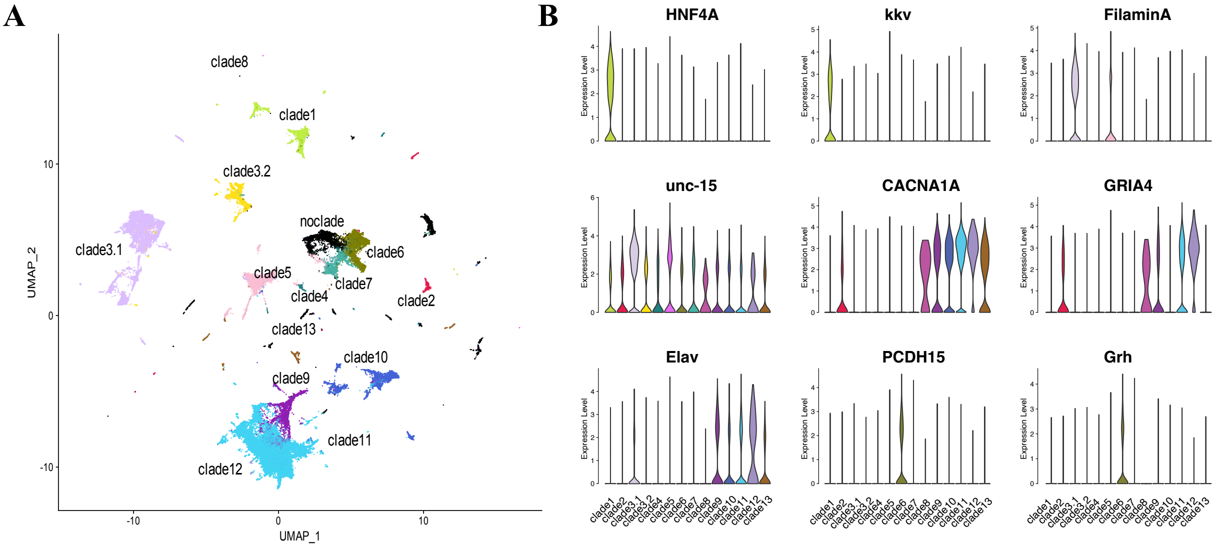
A
B
clade8
clade1
clade3.2
clade3.1
noclade
clade6
clade5
clade7
clade4
clade2
clade13
clade10
clade9
clade11
clade12
UMAP_1
UMAP_2
HNF4A
kkv
FilaminA
unc-15
CACNA1A
GRIA4
Elav
PCDH15
Grh
Expression Level

| Gene | Genome v2.1 ID | Probe sequence | Imaging method | Comments |
|---|---|---|---|---|
| Prox | XLOC_021539 | [illegible] | Confocal | |
| ROpsin | XLOC_028295 | [illegible] | Confocal | |
| Pkd1 | XLOC_027897 | [illegible] | Confocal | |
| Ptf1 | XLOC_003959 | [illegible] | Confocal | |
| COpsin1 | AY692353.1 | [illegible] | Confocal | Not from v2.1 Genome. Has previous accession number published on NCBI. |
| Emx | XLOC_069317 | [illegible] | Confocal | |
| Tll | XLOC_002678 | [illegible] | Confocal | |
| OTX | XLOC_060103 | [illegible] | Confocal | |
| Arx | XLOC_044659 | [illegible] | Confocal | |
| NCALD | XLOC_040379 | [illegible] | Confocal | |
| C-Opsin1 Intron2 | XLOC_012133 | [illegible] | Confocal | |
| Brn3 | XLOC_000129 | [illegible] | Confocal | |
| StMhc | XLOC_045336 | [illegible] | Confocal | |
| Dbx | XLOC_010144 | [illegible] | Confocal | |
| Msx | XLOC_050016 | [illegible] | Confocal | |
| Vglut | XLOC_008985 | [illegible] | Confocal | |
| Phc2 | XLOC_011941 | [illegible] | Confocal | |
| Go-Opsin1 | XLOC_005894 | [illegible] | Confocal | |
| FoxQ2 | XLOC_019990 | [illegible] | Confocal | |
| Bsx | XLOC_045810 | [illegible] | Confocal | Probe sequence deviates significantly from published gene transcripts. |
| Hoxa1 | XLOC_037852 | [illegible] | Confocal | |
| ER81 | XLOC_011095 | [illegible] | Confocal | |
| Six4 | XLOC_059161 | [illegible] | Confocal | |
| Neuropsin2 | XLOC_026826 | [illegible] | Confocal | |
| Fezf | XLOC_021517 | [illegible] | Confocal | |
| CoE | XLOC_028320 | [illegible] | Confocal | |
| GATA123 | XLOC_013055 | [illegible] | Confocal | |
| Bmal | XLOC_009283 | [illegible] | Spinning Disk | |
| FunkyOpsin | XLOC_006912 / XLOC_006910 | [illegible] | Spinning Disk | |
| Itpr3 | XLOC_005874 | [illegible] | Spinning Disk | |
| Ptf1 | XLOC_003959 | [illegible] | Spinning Disk | |
| R-Opsin2 | XLOC_014658 | [illegible] | Spinning Disk | |
| R-Opsin5 | XLOC_027716 | [illegible] | Spinning Disk | |
| Robo1 | XLOC_046455 | [illegible] | Spinning Disk | |

| | | | |
|---|---|---|---|
| StMhc | XLOC_045336 | AUgUCAUACgACgCAUCAgACCCAgAUUACCAgUAUCUggCCgUggACAAgAAggCCUUgCUCAAgAAgCAggCUgAAgCCggAgACACCAAgAAgAUgUgCUUCgUCCCCgACgAgAAggACggAUggAUgAAgggAgAgAUCCAgAgCACCAAgggAgACgACgUCgUCgUCCUCAUggACAAAggAggAgAggUCACAgUgAAgAAggACAAggUCCAgCAgAUgAACCCUCCCAAgUUCgAgAUgAUUgAAgACAUggCCAACUUgACCUACCUCAACgAggCCUCCAUUUUgCACAACUUgCgUACUCgCUACUACAACUCUUUgAUCUACACAUACUCUggCCUUUUCUgCgUggCCAUCAACCCCUUCAgAAgACUgCCCAUCUACggCAUgAACgUCgUACUCAAgUUCAAgggCAAgAAgAggAACgAggUCCCACCCCACUUgUUCUgCAUUgCUgACACAgCCUACUCCAACAUgUUgAUggAUCgUgAgAACCAAUCUAUgCUUAUCACUggCgAgUCUggAgCCggUAAgACAgAgAACACgAAAAAggUAAUCUCCUACUUUgCCCAggUggCUgCUUCCUCCgCUggCAAgAAggAggAgAAAgCUgCUgACgACAAgAAgggCACCCUUgAggAUCAgAUCgUgCAggCCAACCCUgUACUUgAggCCUAUggCAAUgCCAAgACCACCAggAACAACAACUCCUCUCgUUUCggAAAgUUCAUCCgUAUUCAUUUUggCCCAUCUggCAAgAUCgCUggUUgCgAUAUUgAgACUUAUCUgCUggAAAAgUCCCgCgUCACAUACCAgCAgCCUggUgUAgAgCgUAACUACCAUAUCUUUUACUggCUgCUCUCUgAUgCCUUCCCAgACUACCAUgACAAACUggUggUgACACCCgACCCCggCCUCUUCUUCUUCAUCAACCAAggCUgCUUgACUgUUgACggAAUggACgAUCCAgAggAAAUgAAACUgACAgACgACgCCUUUgCUgUgUUgggUUUCACCACUgAggAgAAgAUgggUCUgUACAAgUgCACCUgCgCCAUCCUCCACAUgggAgAgAUgAAgUUCAAgCAgAgACCCAgAgAggAgCAggCUgAggCCgAUggAACCgCCgAggCUgAgAAggUCUCCUUCCUCAUgggAgUCAACgCAUCUgACAUgCUCAAgUgCUUgCUCAAgCCCAAggUCAAggUCggAAAUgAgUUCgUCACCAAgUCCCAgAACAUgAACCAggUCAACUACgCCgUggCUgCCCUggCCAAgUCCCUCUACgACAgAAUgUUCAAgUggCUCgUAgCCCgUgUCAACUUgACACUCgACACCAAgAACAAgAggCAgUACUUCAUCggUgUgUUggAUAUCgCCggAUUCgAgAUCUUCgAgUACAACACCUUUgAgCAgCUUUgCAUCAACUACACCAACgAgCgUCUgCAgCAgUUCUUCAACCAUCACAUgUUCgUgCUggAgCAggAAgAgUACAAgAAggAgggCAUCCAgUgggAgUUCAUUgACUUCggUAUggAUUUgCAgCAgUgCAUUgACCUgAUUgAgAAgCCCAUgggUAUCCUgUCCAUCCUCgAggAggAgUgCAUgUUCCCCAAggCCUCCgACgACUCCUUCAAggCCAAgAUgUACgAgAACCACUUgggCAAgUCCCCCAACUUCggCAAgCCCAggCCCAACAAgAAUgCAAAgUUCgAggCCCACUUUgAACUgAAgCACUACgCCggAACUgUCggCUACAACAUCUCUggAUggCUggACAAgAACAAggAUCCCAUCAACgAgACUgUggUgACCCUgCUgUCCCAgUCCAAggAgCCCAUUgUggCCCACUUgUUCCAggUCCCCAAggAAgAAgCCggCggCggCggUggCAAgAAgAAgAAgggCAAgggAUCUgCCUACCAgACCAUCUCUgCUACCCACAAggAgCAgCUUUUCAAgCUCAUgCACAACUUgCACACAACAUgCCCUCACUUCgUgCgUUgCCUCAUCCCCAAUgAgAUUAAgACUCCAggUgUCAUUgAUgCUAUgCUCgUCCAgAACCAgCUgCAgUgUAACggUgUACUUgAgggUAUCCgUAUUUgCCgUAAAggUUUCCCCAACAggAUgAUCUACUCUgAgUUCAAgCAgAgAUACUCCAUCCUggCUCCCAACgCCAUCCCCCAgUCUggCUUCgUUgACggCAAggUCgCUACUgAgAAggUCCUUggCgCCCUCgACCUCggAgACAACUACCgUCUCggACACACCAAAgUCUUCUUCAAggCUggUAUCCUCggUCAgCUUgAggACAUgCgUgAUgAgCgCUUggCCAAgAUCAUCUCCCUCUUCCAggCCUUCAUCAgAggCUACUUgAUgCgCCgCAgCUACAAgAAgCUCCUCgACCAgCgUACCgCCCUgAgCAUCAUCCAgAggAACAUCCgUCgCUggCUCACAUUgAggAACUggCAAUggUggAAgUUgUACACCCgCgUAAAgCCUCUUCUgUCCAUCgCCCgCCAggAggACgACAUgAAgAAgAAggAggAAgAgUUgAUCgAgACCAAggAgAAgUUggAgAAAUUggAgAAgAUCAAgAAggAgUUggAggAACAgAACgUCgUCCUUCUCCAggCCAAgAACgACCUgUACUUgCAgAUgCAggCUgAgCAggACAACgCUggCgACCAggAAgAgCgCAUUgCCCAgCUgAUCggACAgAAggCUgAgUACgAAgACCgCUUgAAggAACUCgAAgAgCgCCUCggAgAggAggAgggAgCCACCgAAgAgUUggAAgCCAAgAAgAAggCUAUggAAgACAAgAUCggCggACUCAAgAACgACgUgUCCAACUUggAACUCAACUUgCAgAAggCCgAgCAggAgAAgCAgUCCAAggACAACCAgAUCAAgACCCUCAACgACgAgAUggCCCgCCAggACgAgUCCAUCgCCAAgUUgUCCAAggAgAAgAAgAACUUggAAgAgACCAACAAgAAgACCCAggAggAUCUUgCCgCUgAggAggACAAggUCAACCACCUgAACAAgCUCAAgCAgAAgUUggAgCACAACAUCgAUgAgUUggAAgAgAACUUggAgCgUgAgAAgAAgggACgUggCgACUUggACAAggCCAAgAggAAggUCgAgAgCgAgCUCAAgCUCACCCAggAgACCgUUgAggACCUUgAgCgCgUCAAgCgUgACCUCgAggACAACAUCCgCAAgAAggACCAggAgAUCAgCggCAUgAACAgCCgUCUggAggAUgAgUCCAACUUggUUgCCCAgCUCCAgAAgAAgAUCAAggAgCUCAAUgCCCgCAUUgAggAgCUCgAggAggAgCUUgAggCUgAACgCCAggCCAgAAgCAAggUUgAgAAgCAgCgUgCUgAgCUgACCCgUgAAUUggAUgAgCUCUCUgAgCgUCUUgACgAggCUggUggCgCCACUgCUgCCCAgAUCgACUUgAACAAgAAgCgUgAgUCUgAgCUgCAgAAACUgCgCCgUgACUUggAggAggCUCAgAUCCAgAACgAgAgCAACUUgCAggUCAUCAAgAAgAAgCAgCAggAUgCCAUCAACgAgCUgUCCgAACAggUCgACAACCUCACCAAggUCAAgAgCAAgAUUgAgAAggAgAAgAACUCUCUgAAgUCCgAggUUgAUgACCUCCACUCUCAggUUgAgCAUgUCACAAAgAgCAAgUCCUCUgCUgAgAAgCUCgCCAAgCAgUUggAAggCCAgCUUUCCgACgUCAACUCCAAAUUggAUgACUCUggACgCAACAUCCAggACAUCUCCUCCUCCAAgAgCCgCCUCCAgggCgAgAACUCCgACUUgUCCCgCCAgCUCgAggAAgCUgAgAgCCAggUCAACCAgCUCACCAAgCAgAAgCAggCUUUggCCAAgAgCUUggAAgAgACCAAggCUAACCUUgAAgAUgAAUCCCgUggCCgAUCCAAgCUCCAgggUgAggUgCgCAACUUgCAggCCgACUgCgACCAgCUCCgUgAgCAgCUUgAggAggAACAAgAgggACgUggUgAUgUACAgCgUgCCCUUACCAAggCCCAggCUgAggCCggCgAAUggCgCCgCAAgUgCgAgUCCggCgAgggUggAgUCCgCUCUgAggAgUUggAUgACCUCAAgAggAAgAUgAACgCCAAgCUCCAggAUgCCgAAUCUCAgUUggAggCUgCCCUCUCCAAggUCAgCUCCCUCgAgAAgAACAggAACCgCCUCCAgggUgAAgUUgAggAUgUCACCAUCgAAAUggAgAggUgCCAggCCAUUgCCAACCAggCUgAgAAgCgCCAgCgCUCCUUCgACAAgACCAUCgAUgAgUggAAgAAgAAgUCUgCUgACCUCCAgggUgAggUUgAgAACgCCCAgCgUgAAUCCCgUggCAACgCCgCUgAggUCUACAAACUCCgUgCCCAgAUCgACgAgUCUCACgACUCCgUUgAggCCCUCCgCCgUgAgAACAAgAACCUgUCUgAUgAgAUCCACgACAUCACAgAACAgCUCggAgAgggUggCCgCUCUgUCCAUgAAUUggAgAAgAgCCgCAAgCgUUUggAgAUggAgAAggAggAgCUCCAgUCUgCCUUggAggAggCUgAggCUgCCCUUgAACAggAggAggCCAAggUCAUgCgCUCCCAgAUCgAgAUCUCUgCCAUCCgCCAggAgAUUgACAggAggCUCgCUgAgAAggACgAggAgUUCgAgAACACCCgCAAgAACCACCAgCgUgCCCUCgACAgCAUgCAggCUUCCCCUCgAggCCgAgUCCCgUggCAAggCUgAggCCCUgCgCAUgAAgAAgAAgCUCgAgUCUgACAUCAACgAgCUUgAggUUgCCUUggAUggCgCCAACCgUggCCgUgCUgAggCUgAgAAgAACAUCAAgAAAUACCAggCUCAgCUCCgUgAgCUCCAgCAgCACgUUgAggAggAACAgCgUgCCCgUgAUgAggCCCgUgAACAgUACCAggCCUCUgAgCgCCgUgCCAACAUCUUggCUggUgAgAUUgAAgAgUUgAggACCCAgUUggAggCUgCUgAgCgUgCCCgCAAggCCgCUgAgggAgAgCUCCACgAggCUgCCgACCgCgUCUCUgAgCUUUCCACCgCCAACgCCUCUCUCCAggCUgCCAAgAggAAgUUggAgACCgACAUCCAggCCAUgCAgAgCgAUCUUgAggAUCAgUCCAAUgAgCUCAgAUCCgCUgAAgAgCAggCCAAgAAggCCAUggCUgAUgCCgCCAggUUggCUgAggAACUCCgCCAggAACAggAgCACUCUUCUCAgAUUgAAAAgAUgCgCCgCAACCUUgAAUCCCAggUCAAggAgCUCCAggCUCgUCUAgAUgAggCUgAggCUUCCgCCCUUAAgggAggAAAgAggAUgAUCCAgAAAUUggAACAgAgggUgCgUgAgUUggAggUUgAACUCgACAACgAACAgCgCCgCCACggUgAgACCCAgAAgAACCUgCgCAAACAggACCgCCgCCUUAAggAgCUggCCUUCCAgggUgAggAggACCgCAAggCCCAggAgAgACUUACCgACAUgAUUgACAAACUCCAgAACAAgAUCAAgACCUACAAgCgCCAggUCgAggAggCCgAggAAAUCgCUgCUgUUAACCUggCCAAAUACCgCAAggUACAgCACgAgCUUgAggAUgCUgAAgAgCgCgCCgACAUggCUgAgAACACCCUCCAgAAACUgCgUgCCAAgAACCgCAgCUCUgUCUCUgCCACCAggACCACCAgCgCgggUggUgUAACCgUCACCgUCAAAgAgAAACgCACACgCACAUCUgAgAgACUACAACAUgggAAgAUACAACAgCACCCAUggACUCAACCgAUCgUCUUCAAUUCgCggCUCCAAUUCUUCCCUgAAUUUUUCCCUUCgCCCUggCUCAUCCAAUgggggAgACAgUgA | Spinning Disk |
| Tbx6 | XLOC_025705 | ATGCTTTCCCTTCTCCCTCCGACGTCCAGCGGATCCAACATGTTTGAAGAAGACCTTCTGGGGCAGAGAGCCAGGTCCTTTGGGCCCCTGTCACACAGCCTGATGCCCGGACCCCCTCAGGCCTTTCCCTTCACCGGCTTCCCTATGCACCCAGCCCACGACCCCACCATGGCCTTCACAGGTACATGGCAAAGTCAGCAGTTCGGAGGATTCCACATGGGCCACGACCCGAAAAGGATGCTGCCAAGCAGTCAGAGAGAACACAGGGACAAAAATATCAAGATCATCTTGGAAAACAGAGACCTATGGACCAAATTCCACAGCCTGGGGACAGAGATGATCATCACAAAAACTGGCAGACGGATGTTCCCTACATACAAAGTATCCTTGGAGGGCCTGGACCCCCATGCCAAGTACATCTTACTGATGGACATAGTCCCCCTGGACGACTGCAGATACAAGTACCACAACTCCGAATGGGTGGTCACAGGAAAGGCCGAGCCCCATATGCCAGGAAGGCTCTACATCCATCCGGACTCCCCTGCATCCGGAGCACACTGGATGAAGCAGCCTGTCCCCTTCCACAAGCTGAAGCTGACCAATAATAACCTCGACCAAAACGGACATATAATCCTCAACTCCATGCACAAGTACCAGCCCAGAATCCACATCGTGCAGGCCAACGACATATTCTCCATGAGGTGGAATTCATTCAACACTTTTGCTTTTGAAGAAACTGTATTCATCGGAGTGACAGCCTACCAGAATGAACAAATCACACAGCTCAAGATCGACAACAACCCGTTCGCCAAAGGATTCAGGGACAACGGCATGGGCAGAAGATACGAAAACCCCCATAATAATGAAGTGAAGGCCTTAAGAGACCACAGGCTGCATTTGAAAAGAGCCCTGATGCCAGATACACCGAGCAAAGACAACCACGACCATCACGATGACAAGAAAAGATTGAAGACAATGGTCAGTTCGAGCAATCACTCGTCACCGGAAGGAAAGGAAGGACACGGGGGTCACGGAGGATCTTCCGGTCACCAGAGGACTTCCGGCGGAGGGCACATGGACCACGAACTGAAACCATCAGTTGCAGCGGAAGTTGACGATCTCCCTTCCTGTCAGGTCATCAAGGGAAGGTCATCCCTCTCCCAGCAGCAGCAGCCCCTGAGTCGGCCAAGTAATAAGTCCCCCGGCCCCTCCCCTTCCTCCTCAGCCCCGTCAAGCCGGGCCCCCTTGCAGATCTCCCCCCAGTGTCAGTATTTGCCCCAGTGCCCGTCTCCCTACCTCCAGGGCTCCTCTCCCCCCGGATCCGGATGTGACCAGGTCTACTACCCCCACCATCATTCCAACATGGCCATGCAGATGGCCAACTCCGGGCTCCTTCAGCCAGTCACCTCATCCGGATTGCTCCAACCAGTATCAGGCGCCCTCAGCCTCAGCCACATGGGAGCAGGGCAGCAAGTTCAGACATTCCCCGGCCAAAACAGCGCTTGTGCCCTGGCTGCTGGCCCTCCTGGCTCCTCGCAGAGCCCGGCCCCTGGCTACGGGGGCATCTCCGGGGGACAAGGGGCCCACTCCTCCCTCTCCTCCTGCACCTACATGCAGTCGCCCAACCAAACTTACGGACACTTACCATCCTCCATGTCGGTCATGAACATGGCCCCCTCTCACTTTGCTGGTCCCATGGCCTGA | Spinning Disk |

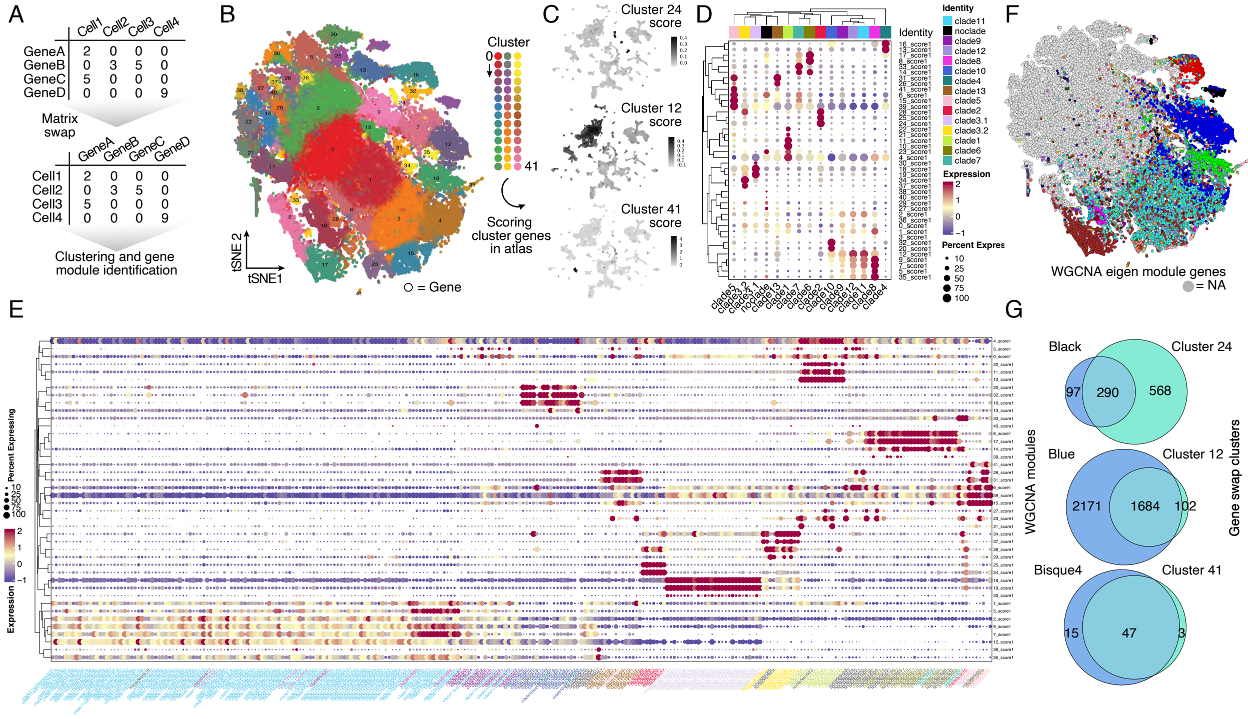
A
Cell1 Cell2 Cell3 Cell4
GeneA 2 0 0 0
GeneB 0 3 5 0
GeneC 5 0 0 0
GeneD 0 0 0 9
Matrix swap
GeneA GeneB GeneC GeneD
Cell1 2 0 0 0
Cell2 0 3 5 0
Cell3 5 0 0 0
Cell4 0 0 0 9
Clustering and gene module identification
B
Cluster
0
41
Scoring cluster genes in atlas
tSNE 2
tSNE1
○ = Gene
C
Cluster 24 score
Cluster 12 score
Cluster 41 score
D
Identity
clade11
noclade
clade9
clade12
clade8
clade10
clade4
clade13
clade5
clade2
clade3.1
clade3.2
clade1
clade6
clade7
Expression
Percent Expres
10
25
50
75
100
F
WGCNA eigen module genes
= NA
E
Percent Expressing
Expression
G
Black
Cluster 24
97
290
568
Blue
Cluster 12
2171
1684
102
Bisque4
Cluster 41
15
47
3
WGCNA modules
Gene swap clusters

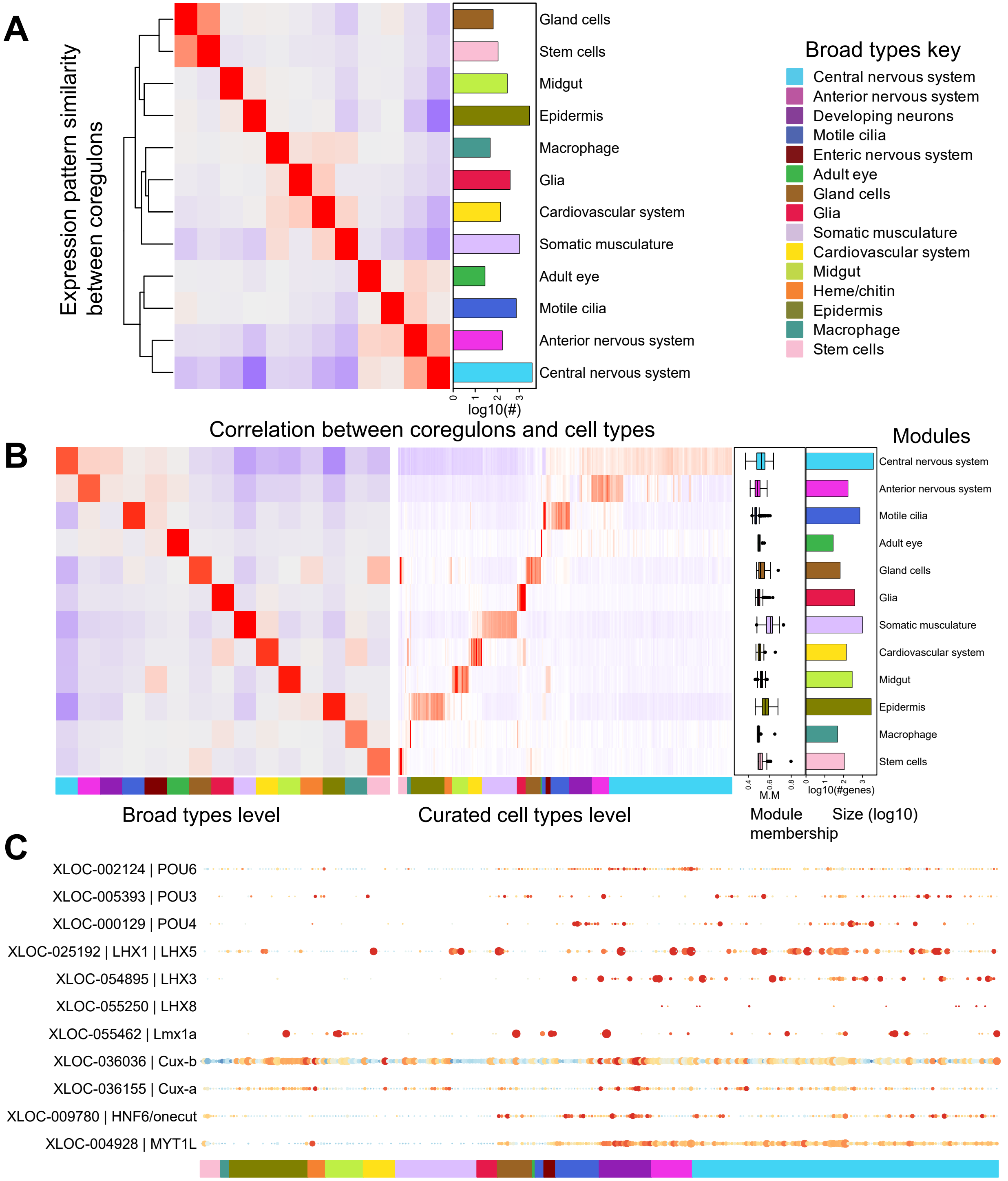

### Supplementary Figure for Figure 3

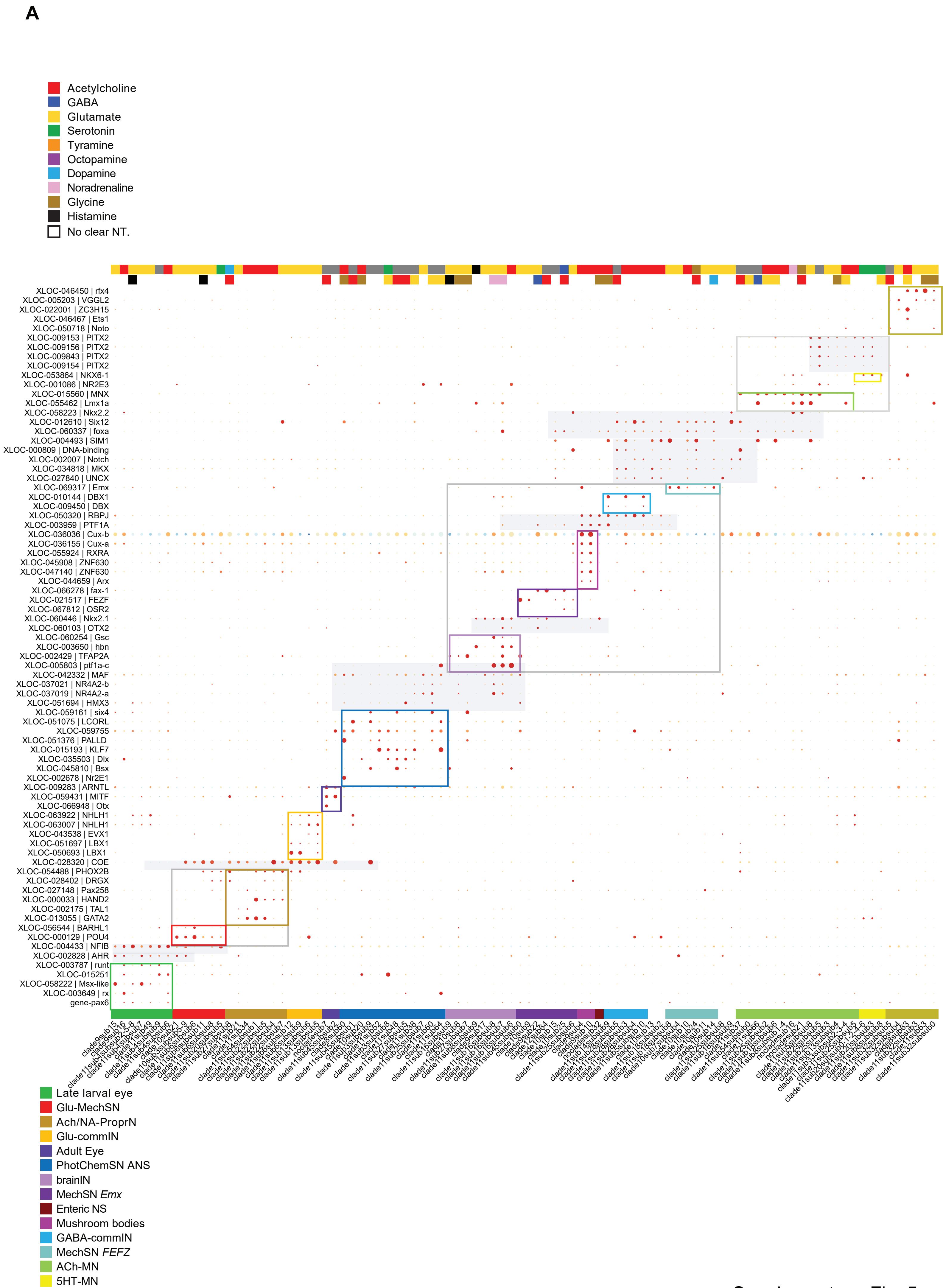

Supplementary. Fig. 5

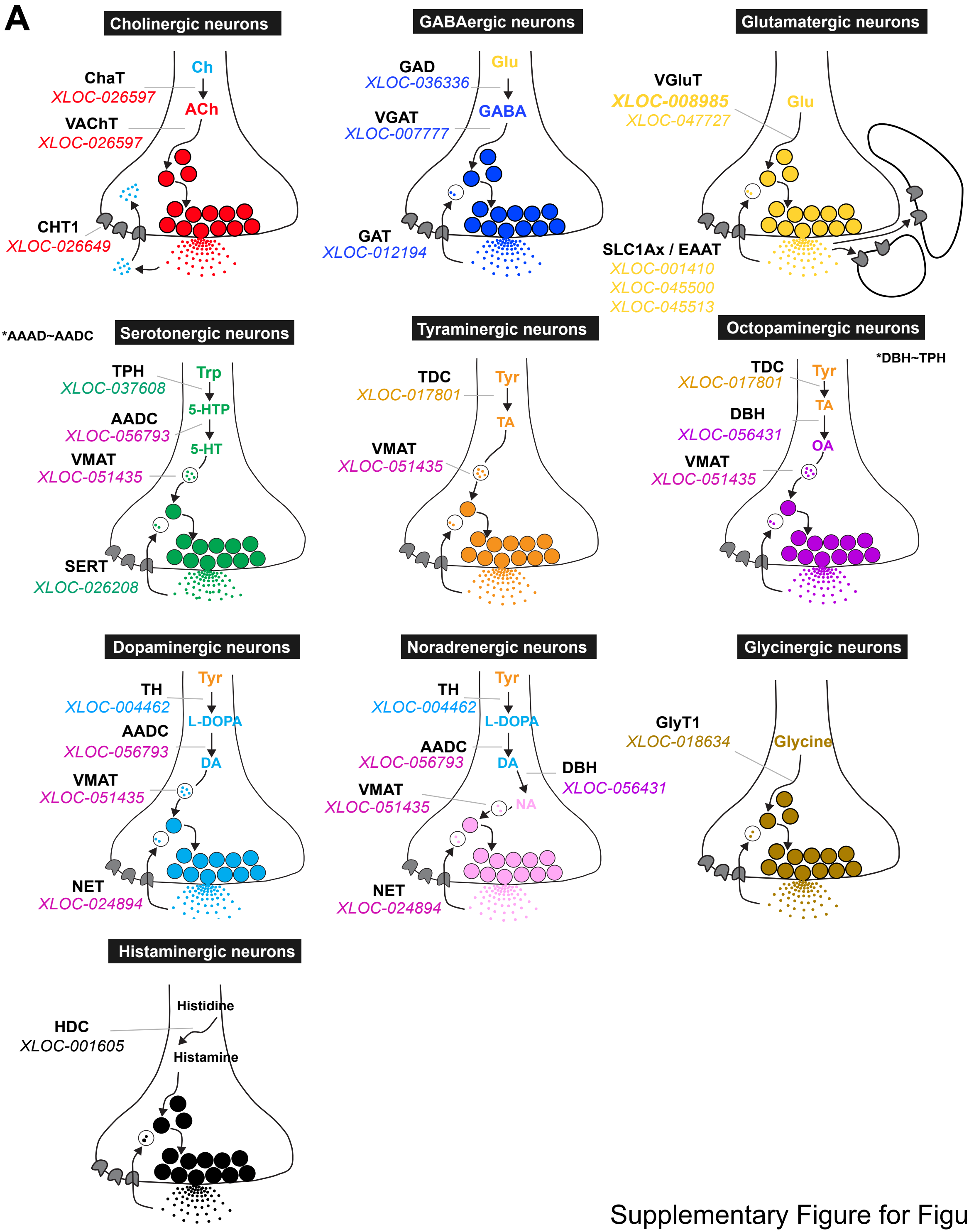

Supplementary Figure for Figure 4

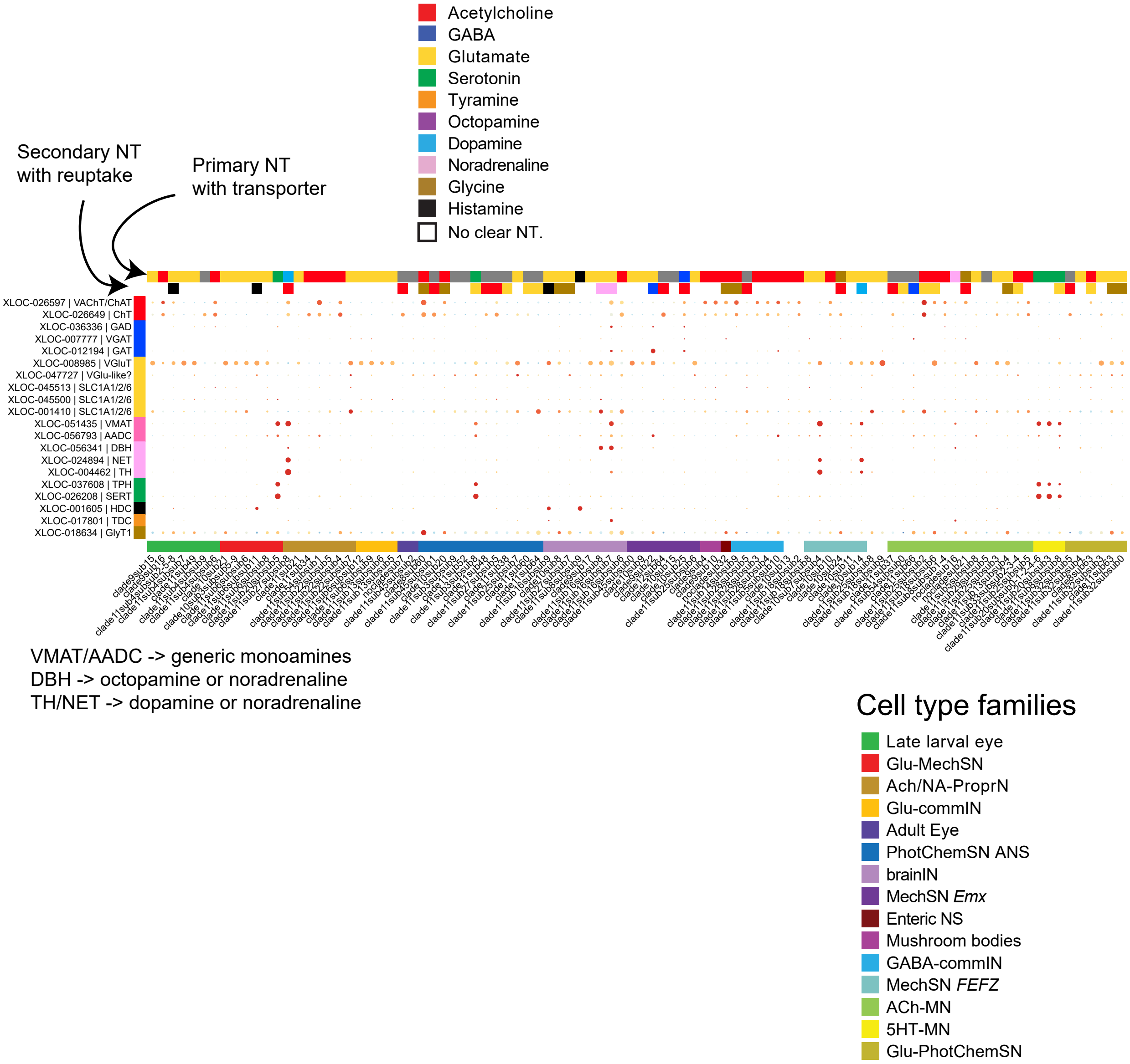

Supp. Figure 7

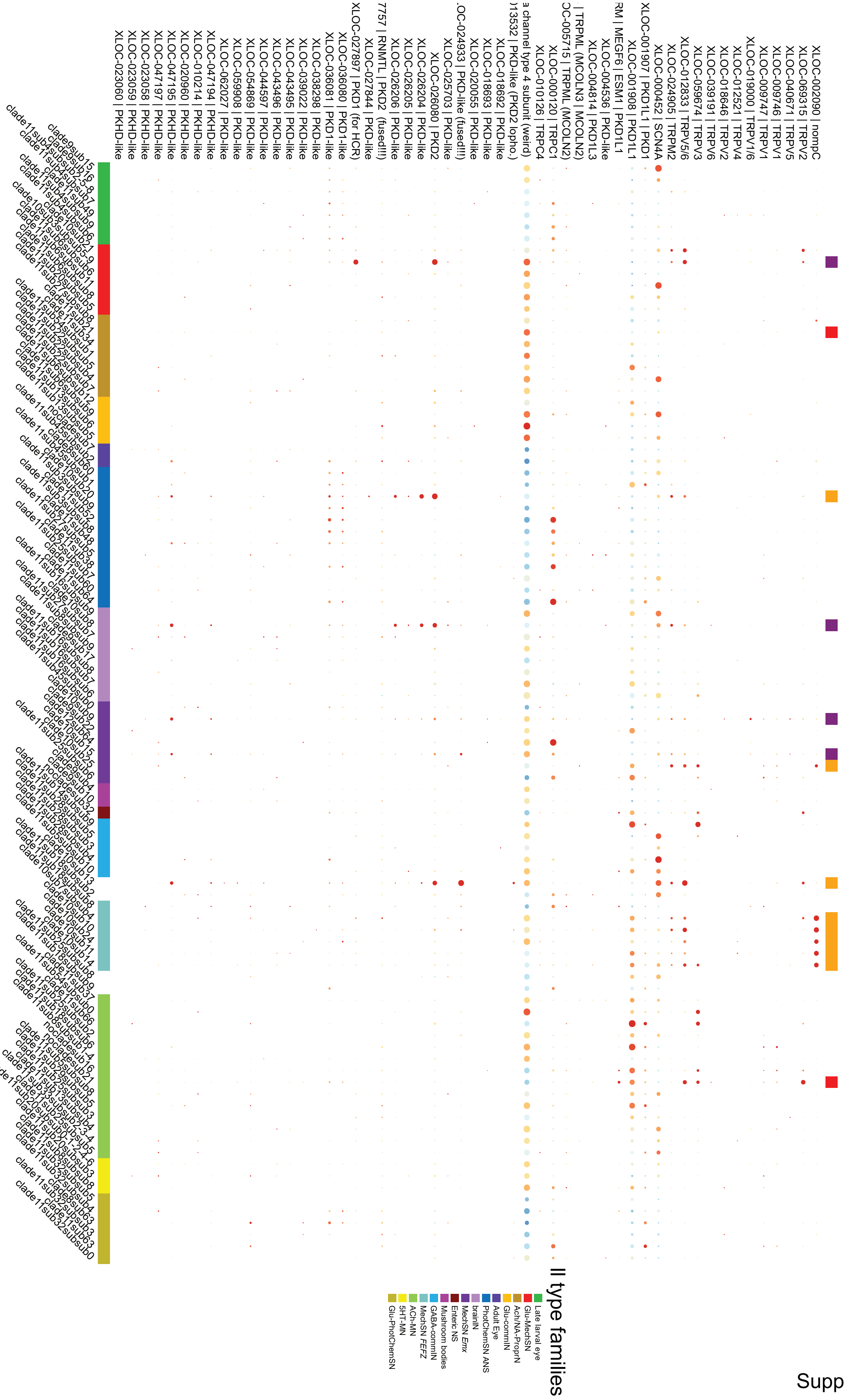

Supp. Figure 8

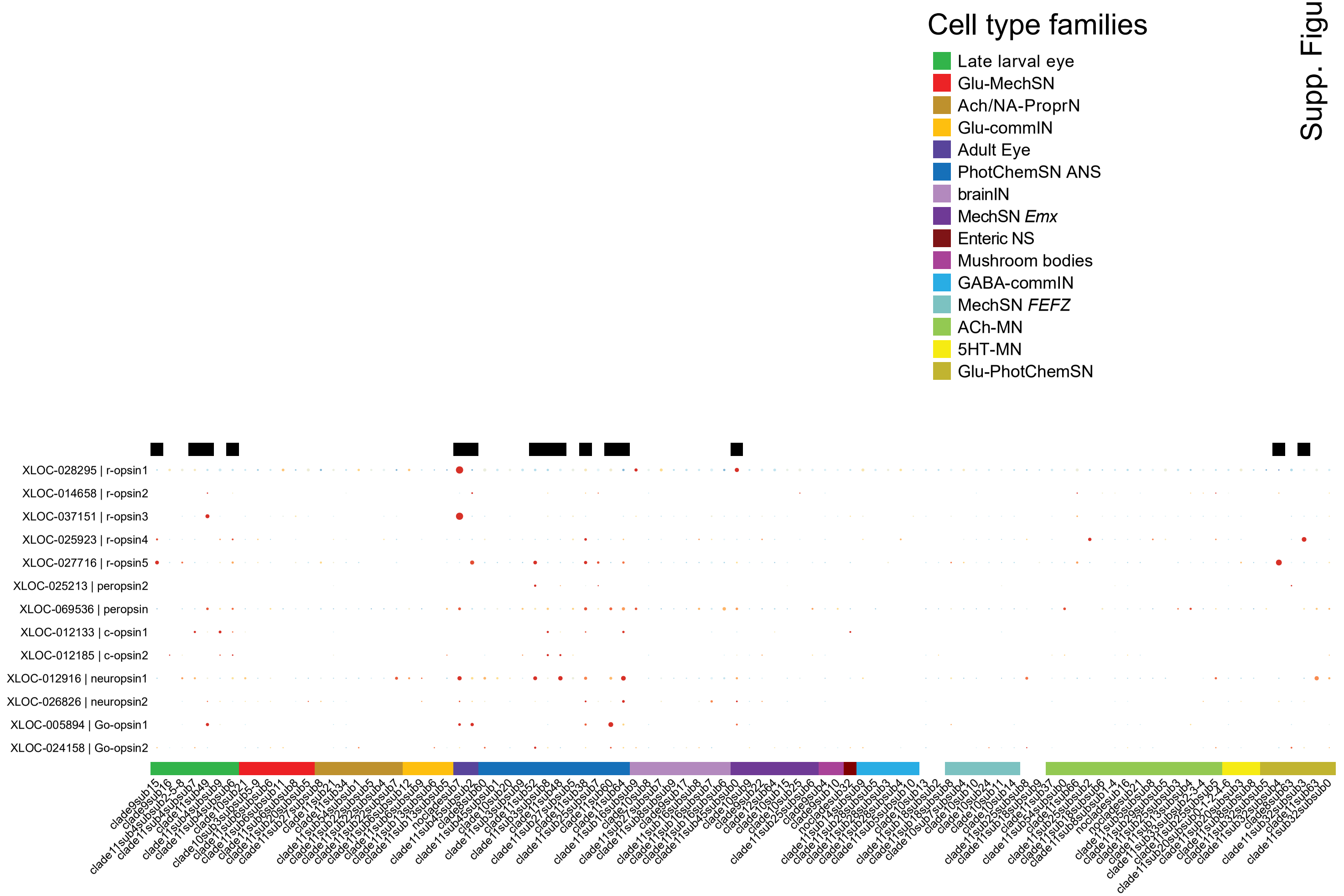

Supp. Figure 9

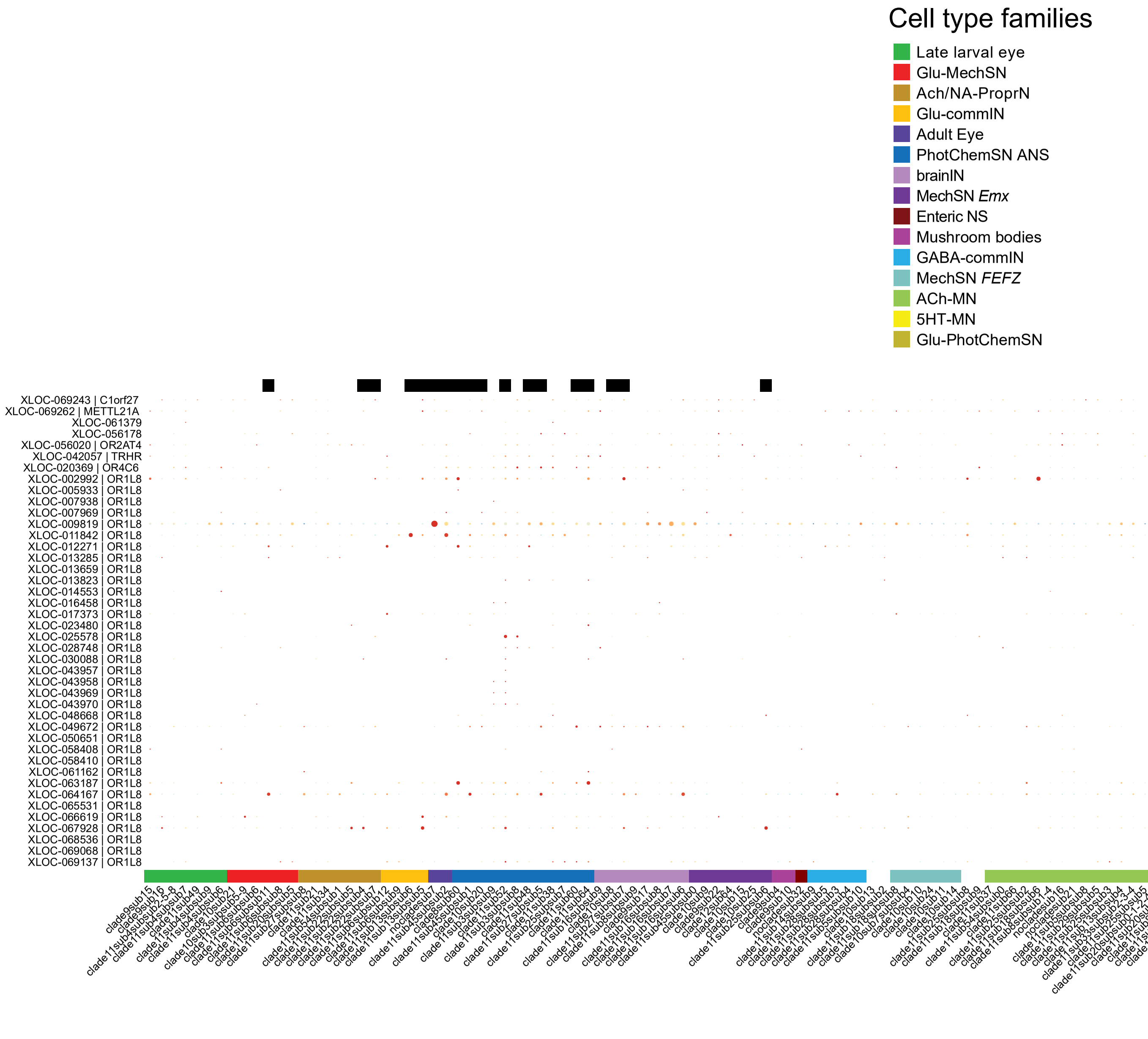

Supp. Figure 10

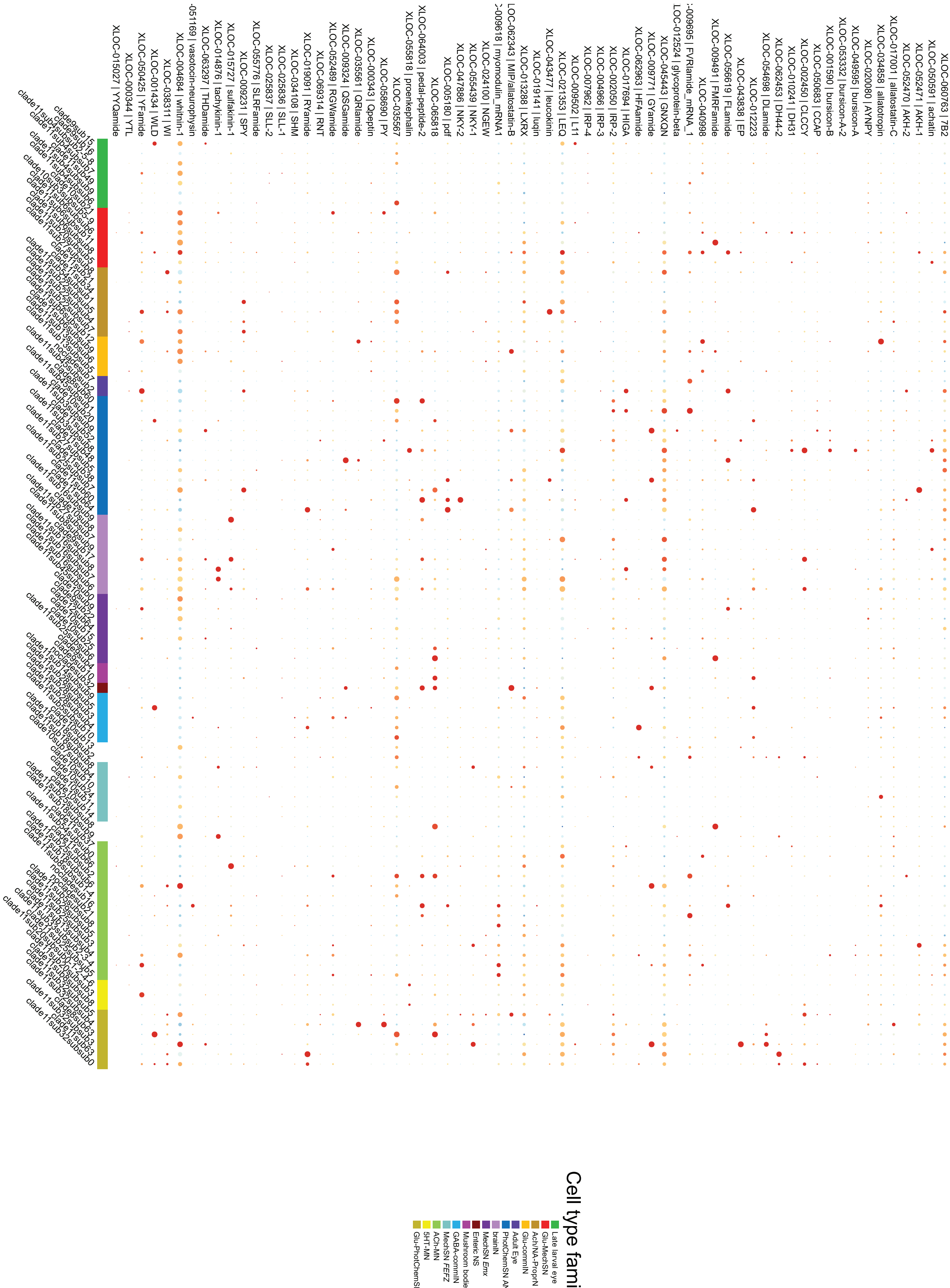

Supp. Figure 11
